# Inhibition of CREB binding and function with a dual-targeting ligand

**DOI:** 10.1101/2023.09.03.556086

**Authors:** Yejun Liu, Stephen T. Joy, Madeleine J. Henley, Joel A. Yates, Sofia D. Merajver, Anna K. Mapp

## Abstract

CBP/p300 is a master transcriptional coactivator, regulating gene activation by interacting with multiple transcriptional activators. Dysregulation of protein-protein interactions (PPIs) between the CBP/p300 KIX domain and its activators is implicated in a number of cancers including breast, leukemia, and colorectal cancer. However, KIX is typically considered ‘undruggable’ because of its shallow binding surfaces lacking both significant topology and promiscuous binding profiles. We previously reported a dual-targeting peptide (MybLL-tide) that inhibits the KIX-Myb interaction with excellent specificity and potency. Here we demonstrate a branched, second-generation analog, CREBLL-tide, that inhibits the KIX-CREB PPI with higher potency and selectivity. Additionally, the best of these CREBLL-tide analogs shows excellent and selective anti-proliferation activity in breast cancer cells. These results indicate CREBLL-tide is an effective tool for assessing the role of KIX-activator interactions in breast cancer and expanding the dual-targeting strategy for inhibiting KIX and other coactivators that contain multiple binding surfaces.

---

Protein-protein interaction (PPI) networks are essential regulators of gene expression.^1, 2^ The activator binding domain (ABD) of coactivators and the transcriptional activation domain (TAD) of activators form structurally dynamic and transient complexes that underpin transcriptional initiation.^2, 3^ The transcriptional coactivators CREB binding protein (CBP) and its paralog p300 feature many domains that control several aspects of transcription, including the KIX domain.^4-7^ The KIX domain, which is one of four CBP/p300 ABDs, regulates gene transcription by interacting with other DNA-associated transcriptional activators such as cAMP response element binding protein (CREB) and MLL; dysregulation of KIX-activator PPIs is frequently observed in cancer.^8, 9^ However, the structurally plastic architecture of the KIX domain has made the development of an effective and selective inhibitors challenging.^10-15^ Especially difficult is the binding surface used by the activators cMyb and CREB, as it is large and topologically challenging.^4, 16-19^ Furthermore, cMyb and CREB engage the surface in orthogonal orientations such that concomitant inhibition of both activators is difficult.^12, 20, 21^

We previously developed a dual-targeting peptide, MybLL-tide, that contained both Myb and MLL TADs and was highly effective at inhibiting at least a subset of KIX PPIs.^22^ Here we report a second-generation dual-site inhibitor, CREBLL-tide, a fusion of CREB and MLL TADs that targets KIX in a distinct orientation. Compared with MybLL-tide, CREBLL-tide and its variants have a higher binding affinity and selectivity for CBP/p300 KIX, as well as improved cellular permeability. In addition, CREBLL-tides interact with endogenous, full-length CBP with significant proteolytic stability. Furthermore, CREBLL-tides reduce the viability of breast cancer cells and inhibits expression of CREB-dependent genes. These results indicate that simultaneously targeting both sites of CBP/p300 KIX is a generalizable strategy to develop potent and selective inhibitors of historically undruggable targets such as CREB.

CREBLL-tide consists of the covalently linked transcriptional activation domains of CREB and MLL (pKID_119-147_ and MLL_2840-2858_) (Figure 1A). The NMR solution structure of MLL-KIX-pKID indicates that the challenge for the CREBLL-tide design arose from the need to connect the two proximal N-termini of the MLL and pKID peptides.^18,19^ Additionally, the MLL-KIX-pKID complex shows considerable flexibility at the N-termini of both peptides (Figure S2, Table S2).^13^ To enable several linkers to be investigated, a convergent synthetic strategy was utilized, in which MLL_2840-2858_ and iodo-linker-pKID_119-147_ were first synthesized independently and then coupled by a displacement reaction using the nucleophilic thiol side chain of C2841 of MLL to form a branched peptide, CREBLL-tide (Figure S1). As illustrated in Table 1, CREBLL-tides with short (β-alanine), medium (6-aminohexanoic acid), and long (AEEAc) linkers were prepared by this synthetic strategy. The purity and mass of the peptides were confirmed by LC-TOF (see SI for full synthetic and characterization details).

**Figure 1.**
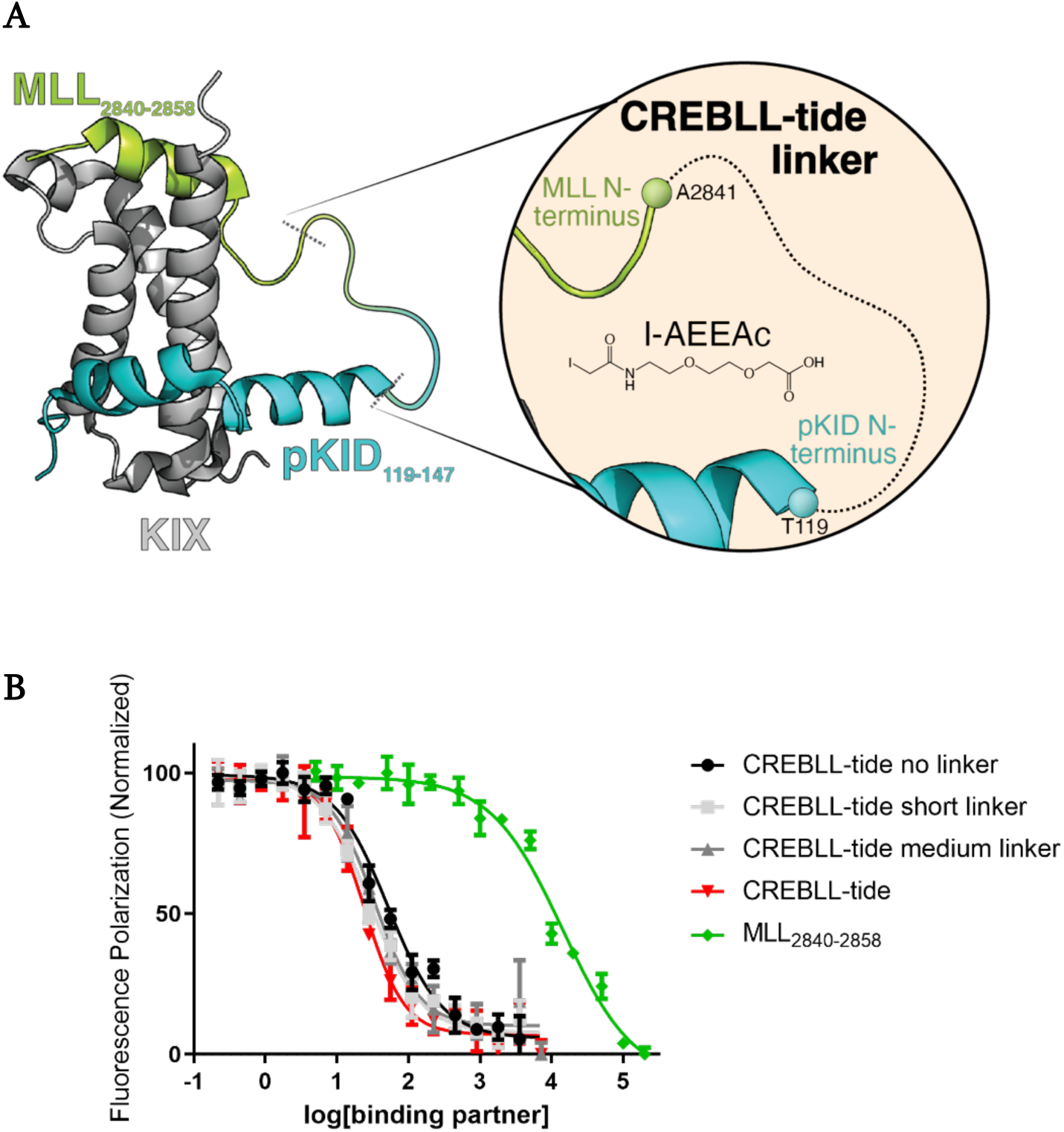
Design of CREBLL-tide and binding affinity of CREBLL-tides with different linker for KIX. (A) Depiction of CREBLL-tide bound to KIX, based upon the NMR structure of the MLL-KIX-pKID ternary complex (PDB: 2LXT). MLL is shown in lime, pKID is shown in cyan. The KIX motif is gray. CREBLL-tide is formed by linking the N-termini of pKID and MLL with an iodo-AEEAc linker. (B) The competitive FP assay of CREBLL-tides with FITC-MybLL-tide for CBP-KIX. The IC_50_ for CREBLL-tide no linker, CREBLL-tide short linker, CREBLL-tide medium linker, CREBLL-tide and MLL_2840-2858_ are 51 ± 1 nM, 29 ± 1 nM, 34 ± 1 nM, 24 ± 1 nM and over 8000 nM respectively. FITC-MybLL-tide is a previously published FITC labeled peptide that binds to KIX with high affinity.^21^ Error bars represent SD from technical triplicates.

**Table 1.** CREBLL-tide constructs used in this study. Schematic of CREBLL-tide constructs consisting of MLL_2840-2858_ and pKID_119-147_ bound by different linkers and labelling groups (R). A tyrosine was added to the N-terminus of MLL_2840-2858_ for determining concentration. FITC = fluorescein isothiocyanate, Ac = acetyl, TAT = Tat-cell penetrating peptide, CLIP6 = CLIP6-cell penetrating peptide, GG = diglycine linker. AEEAc = 8-amino-3,6-dioxaoctanoic acid. Table S1 or Figure S15-S32 for structures.

|  | Linker | R | F2852 | S133 |
| --- | --- | --- | --- | --- |
| CREBLL-tide no linker | / | Ac | F | phos |
| CREBLL-tide short linker | β-Ala | Ac | F | phos |
| CREBLL-tide medium linker | 6-aminohexanoic acid | Ac | F | phos |
| CREBLL-tide | AEEAc | Ac | F | phos |
| FITC-CREBLL-tide | AEEAc | FITC-β-Ala | F | phos |
| FITC-CREBLL-tide-Neg | / | FITC-β-Ala | A | No phos |
| FITC-CLIP6-CREBLL-tide | AEEAc | FITC-β-Ala-CLIP6-GG | F | phos |
| FITC-TAT-CREBLL-tide | AEEAc | FITC-β-Ala-TAT-GG | F | phos |
| CLIP6-CREBLL-tide | AEEAc | Ac-CLIP6-GG | F | phos |
| CLIP6-CREBLL-Neg | / | Ac-CLIP6-GG | A | No phos |
| Biotin-CREBLL-tide | AEEAc | Biotin-AEEAc | F | phos |
| Biotin-CREBLL-Neg | AEEAc | Biotin-AEEAc | A | No phos |

The efficacy of the CREBLL-tides for targeting KIX was evaluated by comparing their ability to inhibit the KIX·MybLL-tide interaction through a competitive fluorescence polarization (FP) assay (Figure 1B). The inhibitory capacity of the CREBLL-tides with different linkers were similar, but the CREBLL-tide with the AEEAc linker showed the best solubility. As a result, the CREBLL-tide composed of MLL_2840-2858_, CREB_119-147_, and an AEEAc linker was chosen for further characterization. A mutant negative control (CREBLL-tide-Neg) was also designed based on this construct in which the linking AEEAc residue was removed, an F2852A installed in the MLL_2840-2858_ construct, and pKID replaced with KID due to the importance of the phosphoserine (pSer133) for KIX-CREB binding (Figure S3).^23^

The binding affinity of CREBLL-tide, pKID_119-147_, and MLL_2840-2858_ to CBP KIX was initially assessed by direct FP experiments. The dissociation constant (K_D_) of the CBP KIX·CREBLL-tide interaction was less than 10 nM, indicating a >40-fold tighter binding than pKID_119-147_ or MLL_2840-2858_ (Figure 2A). A sum of squares error (SSE) analysis was performed to define the upper threshold of the K_D_ for CREBLL-tide indicating a K_D_ ≤ 1 nM (Figure 2B).^24, 25^ In contrast, CREBLL-tide-Neg showed significantly reduced affinity for CBP KIX (K_D_ = 2000 ± 200 nM, Figure S3). To further define the affinity of CREBLL-tide for KIX, a transient kinetics-stopped-flow fluorescence assay was performed.^14, 26, 27^ The results showed CREBLL-tide interacts with CBP KIX with a fast on rate and a slow off rate, and the K_D_ was estimated as 0.32 ± 0.01 nM based on the ratio between k_off_ and k_on_ (Figure 2C, Figure S4, Table S4). In addition, the interaction between CREBLL-tide and CBP-KIX was further tested using an affinity pulldown assay, wherein CREBLL-tide’s ability to bind KIX significantly outperformed biotinylated MLL (Figure S5). Furthermore, CREBLL-tide effectively inhibited the KIX·pKID interaction (K_i_= 98 ± 21 nM) (Figure 2D). Taken together, these results indicate that CREBLL-tide interacts with KIX with high affinity and blocks a key activator PPI.

**Figure 2.**
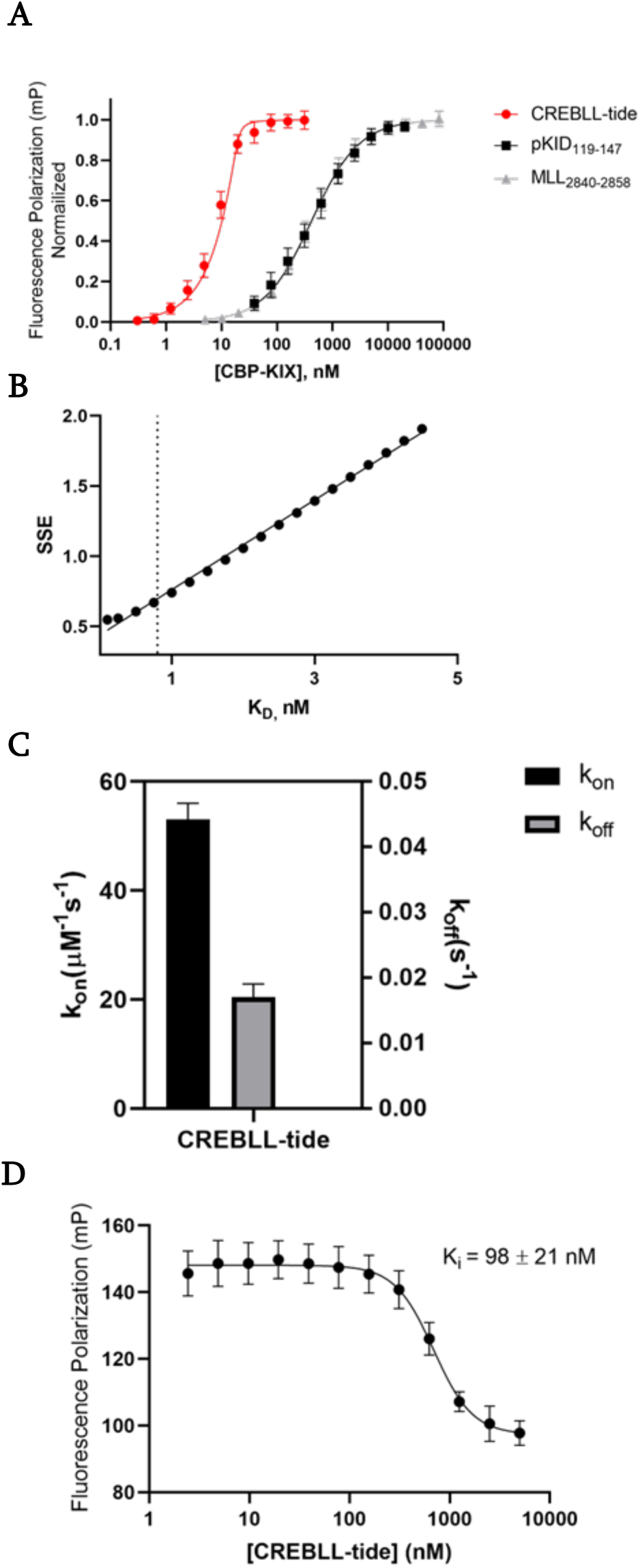
CREBLL-tide interacts with KIX and CBP with high potency. (A) Direct FP binding assay of CREBLL-tide for CBP/p300 KIX. 20 nM of FITC-CREBLL-tide, FITC-MLL_2840-2858_, or FITC-pKID_119-147_ were incubated with CBP-KIX of different concentrations. The dissociation constant (K_D_) of CREBLL-tide was lower than 1 nM, while the K_D_ of pKID and MLL were 410±30 nM and 420±20 nM respectively. Error bars represent SD from technical triplicates of biological triplicates. (B) The sum of squares error analysis of the FP data. The upper threshold for the K_D_ of CREBLL-tide was exhibited by a dashed line, which was determined by calculating the K_D_ corresponding to a 25% increase in SSE above the minimum. The SSE was calculated from three individual experiments composed of technical triplicates. (C) k_on_ and k_off_ for CREBLL-tide based on stopped-flow fluorescence association and 1:1 mixing experiments. Error represents SD from individual triplicate experiments. (D) The competitive FP assay of CREBLL-tide with FITC-pKID_119-147_ for CBP-KIX. Error bars represent SD from technical triplicates of three independent experiments.

To define the binding of CREBLL-tide with full-length CBP in the proteome, affinity pull-down assays were performed using cell lysate from the VARI068 cell line, which is a patient-derived, low-passage triple-negative breast cancer cell line that exhibits activated CREB and overexpressed CBP and p300 (Figure S6).^28^ In addition, VARI068 cells exhibit higher upregulation of CBP and p300 relative to MCF7 and MDA-MB-231 cells (Figure S6). Intact CBP was pulled down with biotinylated CREBLL-tide bound with Neutravidin beads, while no CBP was bound with the biotinylated CREBLL-tide-Neg. Moreover, the affinity pull-down of MybLL-tide with CBP was inhibited by the addition of CREBLL-tide in cell lysate, indicating that CREBLL-tide is capable of interacting with full-length CBP via the KIX domain (Figure 3A, Figure S7, S8).

**Figure 3.**
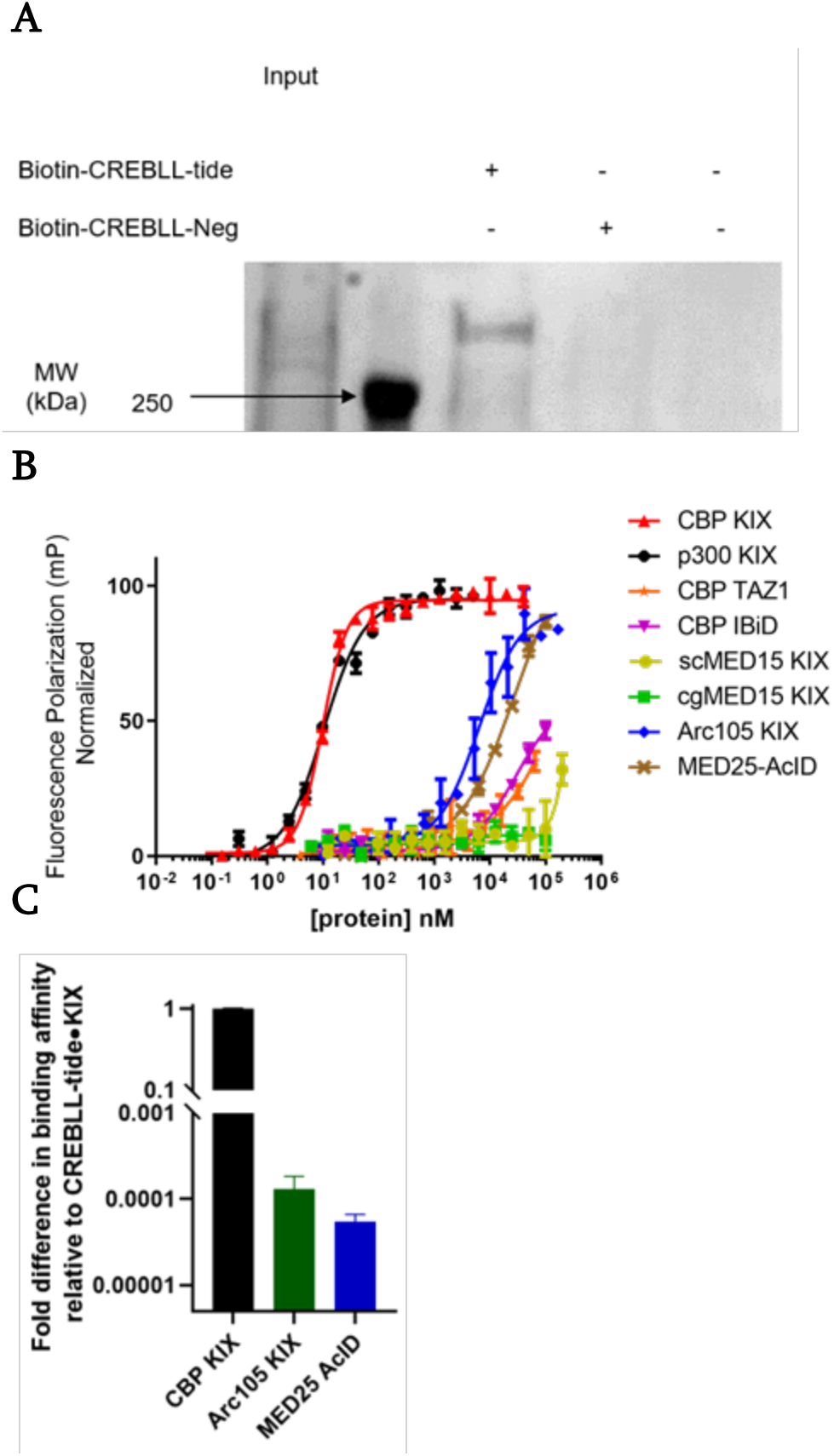
CREBLL-tide interacts with full-length endogenous CBP and CBP/p300 KIX selectively. (A) Western blot for endogenous, full-length CBP and p300 from VARI068 whole cell lysates showing affinity pulldown of CBP via biotinylated CREBLL-tide but not Biotin-CREBLL-Neg. Biotin-CREBLL-Neg was designed based on the weak binding CREBLL-tide-Neg control. 10 μM Biotin-CREBLL-tide or Biotin-CREBLL-Neg were bound to the resin and incubated with the cell lysate for 1 hour. (B) The direct binding FP results of CREBLL-tide vs different proteins. The K_D_ of CREBLL-tide for CBP/p300 KIX, other KIX motifs, other ABDs of CBP and MED25-AcID were shown in SI. Error bars represent SD from technical triplicates of experimental triplicates. (C) The fold difference in the binding affinity of CREBLL-tide to multiple domains relative to CREBLL-tide·KIX interaction. Error bars represent SD from the FP data of (B).

CREBLL-tide was designed to access high binding affinity and more importantly, selectivity, through targeting both binding sites on CBP/p300 KIX. To determine the binding selectivity of CREBLL-tide to KIX, direct FP experiments of CREBLL-tide to different ABDs of coactivators were performed.^22, 28^ CREBLL-tide showed very low binding affinity to other structurally similar but sequentially distinct KIX motifs including cgMED15 KIX, scMED15 KIX, and ARC105 KIX (Figure 3B, Figure S9C-E).^9, 29-31^ Second, other binding domains of CBP (IBiD, TAZ1), which also contain multiple binding sites and are structurally plastic, bound CREBLL-tide very weakly (Figure 3B, Figure S9F, G).^32, 33^ Furthermore, CREBLL-tide did not exhibit binding to MED25 AcID, a coactivator that interacts with similar amphipathic activators (Figure 3B, Figure S9H).^34-37^ Compared to MybLL-tide, CREBLL-tide bound to CBP/p300 KIX with a higher binding affinity and a lower affinity for other ABDs of coactivators such as MED25 ACID and CBP IBiD (Figure 3C). These data demonstrate that, compared with other KIX motifs, binding domains of CBP, or the other ABDs interacting with MLL, CREBLL-tide exhibited at least 8,800-fold higher binding selectivity to CBP/p300 KIX. (Table S5) In addition, the high binding affinity and selectivity of the CREBLL-tide displays the versatility of the dual-targeting strategy for targeting CBP/p300 KIX and provides the foundation for developing additional dual-targeting molecules for inhibiting KIX-activator interactions.

All the previous data demonstrates that CREBLL-tide binds to CBP/p300 KIX with high affinity and selectivity *in vitro*. However, stability and permeability are always challenging for applying peptide-based drugs or tools in cells. To test stability, a human serum degradation assay was performed. After verifying the activity of human serum using an AMC-tagged substrate (Figure S10A, B), CREBLL-tide was incubated with human serum at 37 °C and the solution quenched at different time points. The half-life of CREBLL-tide was measured to be approximately 24 hours. In contrast, the half-life of pKID and KID in human serum was around 6 hours, while the MLL_2840-2858_ peptide was more stable than both CREBLL-tide and pKID (Figure S10C).

To improve the cell-permeability of CREBLL-tide, cell-penetrating peptides (CPP) CLIP6 or TAT with a diglycine spacer were added to the N-terminus of the MLL portion of CREBLL-tide and its negative control mutant.^38, 39^ A FITC tag was attached to the CLIP6-CREBLL-tides and the uptake and localization of peptides were observed by confocal microscopy. CLIP6-CREBLL-tides were incubated with the breast cancer cell lines MDA-MB-231 and VARI068 for 3, 6, and 9 hours, and the MCF7 cell line for 3 hours. CLIP6-CREBLL-tides were detected in both the cytoplasm and the nucleus of different breast cancer cells (Figure 4 A, B and Figures S12-14). Increasing signal of peptide was detected in the nucleus of cells with longer incubation time indicating more CREBLL-tide was transferred into the nucleus as time passed. However, the TAT-linked and CREBLL-tides lacking CPPs were not observed in the nucleus, even after 9 hours incubation (Figure 4A and Figure S11-13). The CLIP6-CREBLL-tides were thus utilized for further studies.

**Figure 4.**
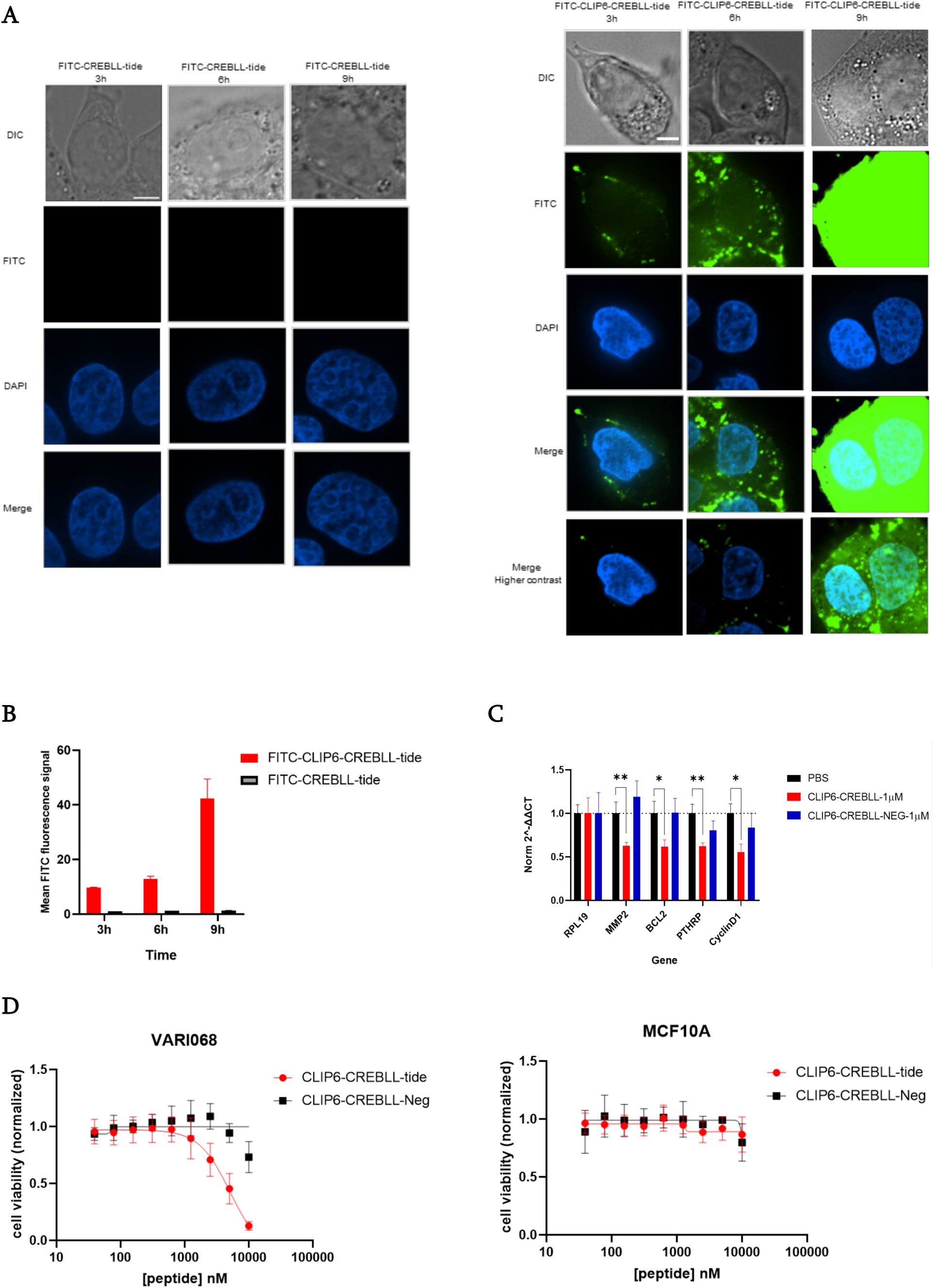
(A) The DIC, FITC and DAPI channel confocal images of VARI068 cells incubating with 10 μM FITC-CREBLL-tide (left) and FITC-CLIP6-CREBLL-tide (right) for 3 h, 6 h, and 9 h. The Merge images are merged by DAPI and FITC channel. Merge images with higher contrast of FITC-CLIP6-CREBLL-tide was also showed. Blue staining in the DAPI and Merge images represent the size and location of the nucleus. Green dots show the location of the FITC labelled peptides in cells. The images are zoomed in to observe the single cell. See Figure S11 for full images. Each scale bar represents 5-micron length in the image. (B) The quantification of the mean fluorescence intensity of the peptides in the nucleus of cells at different times. The mean FITC fluorescence signal within the nucleus was taken and the bar graph represents the average of over 10 cells. Error is the standard deviation from average fluorescence signal of biological triplicates. (C) qRT-PCR analysis of CREB-dependent genes in VARI068 cells after 4 h exposure to 1 μM CLIP6-CREBLL-tide or CLIP6-CREBLL-tide-Neg. RPL19 is the reference gene. Error represents SD calculated from technical triplicates of biological triplicates. **p < 0.01 and *p< 0.1. (D) The VARI068 cell viability (Left) and MCF10A (Right) with the incubation of increasing concentration of CLIP6-CREBLL-tide or CLIP6-CREBLL-tide negative control which lost the binding affinity to KIX. The IC50 of CLIP6-CREBLL-tide for VARI068 was 5.3 ± 8 μM. Error is SD calculated from technical triplicates of biological triplicates.

To verify on-target effects, VARI068 cells were treated with 1 µM CLIP6-CREBLL-tide for four hours and RT-qPCR assays were performed on the CREB-dependent genes including Cyclin D1, PTHrP, MMP2, and BCL-2 which are related to the proliferation, migration, and invasion of cancer cells.^40-45^ These CREB-dependent genes were significantly down-regulated with 1 µM CLIP6-CREBLL-tide after 4 hours. Additionally, the CLIP6-CREBLL-Neg showed moderate or no effect on the transcription level of these genes (Figure 4C). These results indicate that the inhibition of CBP/p300 KIX•activators interaction by the dual-targeting peptide regulates gene transcription related to CBP/p300 KIX in cancer cells.

To test the activity of CLIP6-CREBLL-tide on cell survival, an MTT cell viability assay was performed. After treating VARI068 cells with increasing concentrations of CLIP6-CREBLL-tide, cell viability was dramatically decreased. In contrast, the negative control treatment had only modest effects on cell viability, indicating cell death was not due to the CLIP6 or the CREBLL-tide architecture itself but was likely due to the inhibition of CBP/p300 KIX (Figure 4D). In addition, to detect the toxicity and off-target effects of CLIP6-CREBLL-tide in non-cancerous cells, the MCF10A cell line, which is a non-tumorigenic breast epithelial cell line and not dependent on CBP/p300 and CREB, was used (Figure S14). MCF10A cells were treated with CLIP6-CREBLL-tide and an MTT cell viability assay was performed. The proliferation rate of MCF10A cells did not change significantly after the same treatments indicating cancer cells are more sensitive to the inhibition of the interaction within CBP/p300 KIX by CLIP6-CREBLL than non-cancerous cells (Figure 4D).

The picomolar affinity, high selectivity for CBP/p300 KIX, long-term stability in human serum, excellent cell permeability, and on-target effects on cell viability in breast cancer models demonstrate CREBLL-tide and its derivatives as bona fide inhibitors of CBP/p300 KIX PPI and confirms the feasibility of utilizing dual-targeting peptides in cell models. The results indicate that the inhibition of CBP/p300 KIX leads to down-regulation of CREB-related genes. In addition, the success of CREBLL-tide indicates that branched synthetic can serve as an excellent probe to detect additional functions of CBP/p300 KIX, expanding the architectural possibilities beyond our earlier study of a dual-targeting peptide.^22^

Further studies will focus on differentiating the genes regulated by KIX·activators interaction and the genes controlled by other coactivator-related interactions. Finally, CREBLL-tide’s preferential binding to KIX over other CBP domains represents a unique opportunity to explore how inhibition and rigidification of one domain in such a large, multidomain protein may impact the global structure and capabilities of master coactivators, and we hope to probe broader structural impacts in future studies.

## Supporting information

Supporting information, including methods and data

## Supporting Information

Additional experimental details, materials, and methods of each experiment, peptide sequences, Western bolts, confocal images and LC-MS spectrum.

## Acknowledgements

The authors acknowledge financial support from the National Institutes of Health (GM136356 to A.K.M.; NIH-P30CA046592 to the Rogel Cancer Center Core) and the Breast Cancer Research Foundation (SDM). The authors thank Prof. Wenjing Wang’s lab members past and present for their assistance of confocal microscopy, and helpful conversations. Special thanks are extended to Prof. Wenjing Wang and Jiaqi Shen for sharing their experience in analyzing the results of confocal microscopy.

## Accession Codes

CBP (UniProtKB Q92793)

P300 (UniProtKB Q09472)

CREB (UniProtKB P16220)

MLL (UniProtKB Q03164)

## Table of contents figure

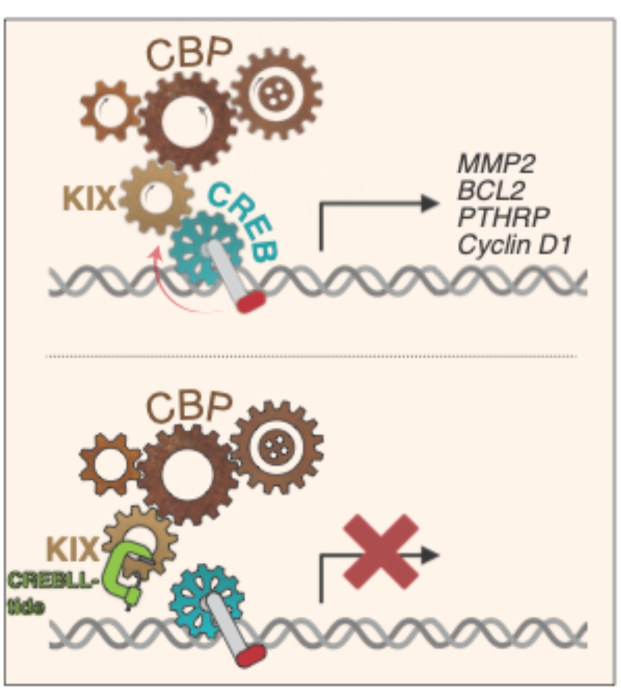

