## Supporting information, including methods and data for "Inhibition of CREB binding and function with a dual-targeting ligand"

### Inhibiting CREB binding and function with a dual-targeting peptide

**Abbreviations.**

| | |
|---|---|
| ABD | Activator-Binding domain |
| AcID | Activator Interacting Domain |
| AEEAc | 8-amino-3,6-dioxaoctanoic acid |
| CBP | CREB-Binding Protein |
| cgMED15 | *Candida glabrata* MED15 |
| CPP | Cell-Penetrating Peptide |
| CREB | cAMP Responsive Element Binding Protein |
| DAPI | 4',6-diamidino-2-phenylindole |
| DIPEA | Diisopropylethylamine |
| DTT | Dithiothreitol |
| FITC | fluorescein isothiocyanate |
| HPLC | high-performance liquid chromatography |
| IBiD | Interferon-Binding Domain |
| KIX | Kinase-inducible domain Interacting domain |
| MLL | Mixed-lineage leukemia |
| MMP | matrix metalloproteinases |
| NMR | nuclear magnetic resonance |
| pKID | Phosphorylated KIX interacting domain |
| qPCR | quantitative Reverse Transcription-Polymerase Chain Reaction |
| scMED15 | *Saccharomyces cerevisiae* MED15 |
| TAD | Transcriptional Activation Domain |
| TAZ1 | Transcription Adaptor putative Zinc finger 1 domain |
| 6-Ahx | 6-aminohexanoic acid |

**General**

Peptide synthesis reagents including trifluoroacetic acid and piperidine were purchased from Sigma Aldrich. Acetonitrile and other solvents were acquired from Fisher Scientific. All primers were acquired from Sigma. All antibodies were purchased from Abcam or Santa Cruz Biotechnology. PowerUp SYBR Green Master Mix for qPCR was purchased from Applied Biosystems and iScript Reverse Transcriptase was acquired from Bio-Rad.

#### Methods

**Peptide Synthesis.** All peptides including CREBLL-tides and their mutants, MLL, pKID, and the CBP IBiD domain (2063-2111), were synthesized by standard Fmoc solid phase peptide synthesis method on a Liberty Blue synthesizer as previously reported.[1] The FITC-tag was added at the conclusion of the synthesis with the incubation of 1.5 eq FITC (ThermoFisher) in 5% DIPEA (Sigma Aldrich) in DMF for 16 hr. MLL-based peptides were cleaved from resin using a solution of 90% trifluoroacetic acid (Sigma Aldrich), 5% thioanisole (Sigma Aldrich), 3% ethanedithiol (Sigma Aldrich), and 2% anisole (Sigma Aldrich) and incubated for 4 hours. pKID-based peptides were cleaved by a solution consisting of 95% trifluoroacetic acid, 2.5% triisopropyl silane (Sigma Aldrich), and 2.5% water and incubated for 4 hours. The cleaved mixture was filtered, and the filtrate concentrated under nitrogen flow. The crude peptides were pelleted by the addition of ice-cold ether. The pellet was dissolved in 20% acetonitrile in water. The solution was frozen in liquid nitrogen and lyophilized to dryness. The pellet was dissolved in 1:1 solution of 0.1% TFA in water and acetonitrile with minimal 50 mM TRIS (pH 8.0). The solution was filtered before being purified by HPLC on a C18 column. Peptide-containing fractions were pooled and lyophilized to a powder. The purity and identity of the peptides was validated by analytical HPLC and mass spectrometry (Agilent 6545 LC/Q-TOF). The CREBLL-tide analogs were synthesized by the incubation of purified MLL and I-Linker-pKID peptides in 50 mM TRIS (pH 8.0) buffer for 4 hours (Figure S1). The mixture was further purified via HPLC, and the resulting purified peptides were collected and validated by Agilent 6230 LC/TOF mass spec.

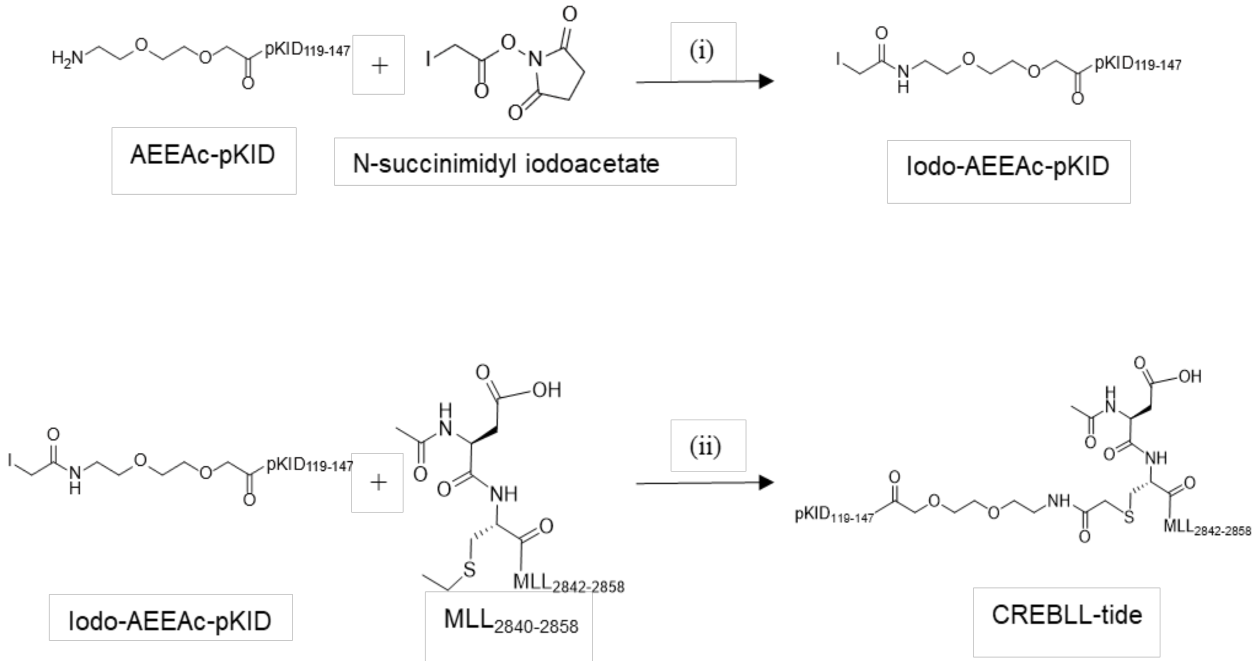


**Figure S1. Coupling of $MLL_{2840\text{-}2858}$ and iodo-$pKID_{119\text{-}147}$.** AEEAc-$pKID_{119\text{-}147}$ reacts with N-succinimidyl iodoacetate to form Iodo-AEEAc-$pKID_{119\text{-}147}$. (i) 0.125 mmol AEEAc-pKID and 0.0375 mmol N-succinimidyl iodoacetate were incubated in 0.5 mL 50 mM TRIS buffer (pH=8.0)for 30 min at room temperature. (ii) 20 μL of 1 mM Iodo-AEEAc-$pKID_{119\text{-}147}$ and 10 μL of 1 mM $MLL_{2840\text{-}2858}$ were incubated in 0.5 mL 50 mM TRIS buffer (pH=8.0) for 4 hours at room temperature. The yield of CREBLL-tide is approximately 40%.

**Protein expression and purification.** CBP KIX, p300 KIX, MED25-AcID and other proteins used in this work were expressed using previously reported methods.[2–7]

**Fluorescence Polarization.** Polarization experiments were performed using previously reported methods.[1,8,9] For direct binding assays, the curves were fit to the observed polarization values as a function of the protein to obtain the apparent dissociation constant,

$$Y = \frac{20 + X + K_D - \sqrt{(20 + X + K_D)^2 - 80X}}{40}$$

"X" is the total concentration of protein. and "Y" is the observed binding percentage at a given protein concentration.

For competitive binding experiments, the curves were fitted to a built-in equation of GraphPad Prism which is "[Inhibitor] vs. response -- Variable slope" to get the $IC_{50}$. The $IC_{50}$ values were converted to Ki values using the apparent Kd value calculated from the direct binding experiments using a Ki calculator.[10] Reported Ki values are the average of three independent replicates with the indicated error representing the standard deviation of the three triplicates.

**Fluorescence stopped-flow affinity assay.** The experiments were performed in the KIX Binding Buffer consisting of 10 mM sodium phosphate (pH = 6.8), 100 mM sodium chloride, 10 % glycerol, and 0.01% NP-40 with 10 mM DTT. The FITC signal was excited at 488 nm and measured at wavelengths >510 nm using a long-pass filter (Corion). The $k_{on}$ was determined by the concentration dependence experiments which were performed by mixing FITC-CREBLL-tide of constant concentration (0.025 μM) with variable concentrations of KIX. The $k_{off}$ value was determined by the low concentration 1:1 CREBLL-KIX association experiments where 5 nM KIX was rapidly mixed with 5 nM FITC-CREBLL-tide. 10 to 15 traces were collected before the final fitting. The data was analyzed using the previously reported procedure.[11,12]

**Cell culture.** MCF-7, MCF10A, and MDA-MB-231 cells were purchased from ATCC and used in accordance with ATCC protocols. VARI068 cells were handled as previously described.[9] MCF10A, MCF-7 and MDA-MB-231 cells were cultured following the protocols of ATCC. All cell lines were grown at 37 °C and 5% $CO_2$.

**Affinity Pulldown assay**. NeutrAvidin Agarose Resin (Thermo Scientific 29200) was spun down at 3000 g for 2 min, washed twice with Biotin Binding Buffer (100 mM sodium phosphate, 150 mM sodium chloride, pH = 7.2), and resuspended in 345 μL Biotin binding buffer. Ten nanomoles of biotinylated peptides were dissolved in 0.1 mL Biotin Binding Buffer. The resin, peptides, and 10 mM DTT were combined in a total volume of 0.5 mL and the mixture was agitated for 1 hour at RT. The resin was spun down at 3000g for 2 min, resuspended in Superblock blocking buffer and agitated for 1 hour at RT. The resin was washed three times using KIX binding buffer (10 mM sodium phosphate, 100 mM sodium chloride, 10% glycerol, 0.01% NP-40, pH =6.8). During the agitation, the breast cancer cells were collected (1 million cells per sample) and washed with cold PBS 3 times. Cells were lysed in cold TRIS lysis buffer (50 mM TRIS (pH = 7.4), 150 mM NaCl, 0.5 mM EDTA, 0.5% TRITON-

X and 10% glycerol) with protease inhibitors (500 μM 4-benzenesulfonyl fluoride hydrochloride, 10 μM Bestatin, 100 μM Leupeptin and 1 μM Pepstatin). The suspension was vortexed vigorously for 10s, incubate in ice for 10 min, then spin down at 14000 g for 30 min. The supernatant was transferred to another tube and spin down again at 14000 g for 15 min. The cell lysate was collected and combined with the resin bound by peptides or unbound. The mixture was agitated for 1 h at 4 ˚C and washed twice. The components bound to the resin were removed by boiling in 5:2:13 Laemmli buffer: β-mercaptoethanol: 0.1 M citric acid (pH = 2.1) before separation on a 4-20% PAGE bis-tris gel. After running at 150 V for 2 h, the gel was transferred to a PVDF membrane using a Trans-blot Turbo Transfer system. The membrane was blocked for 1 hour before incubating with the anti-CBP (sc-7300 from Santa Cruz) and anti-mouse IgG secondary (sc-516102 from Santa Cruz).

**Confocal Microscopy.** Cells were counted and resuspended in 15 mm glass bottom plates (200,000 cells per well). Cells were grown in the incubator overnight. The medium was changed to 1% FBS DMEM, FITC-labeled peptides or DMSO were added to the plates and the plates were incubated at 37 ˚C for different time periods. Cells were washed three times with PBS and fixed in paraformaldehyde and nuclei were stained by DAPI. The cells were washed again with PBS before being imaged using a NIKON TI2-FP confocal microscope. All images were shown with the same contrast except the "Merge Higher construst" images of Figure 4A. The images and the quantification of the fluorescence signal were analyzed using Image J. The images were transferred to 8-bit images and DAPI staining was used to determine nuclear position, shape, and size and assess co-localization with FITC-labeled peptides. The background signal was subtracted before measuring the area integrated intensity. The mean intensity of cells was calculated based on the data from over 10 cells.

**Human serum stability assay.** Peptides were dissolved in 500 mM TRIS, pH 8.0 to a concentration of 500 μM. Peptide solutions and 50% human serum in DMEM were incubated separately in a 37 ˚C metal bead bath for 30 min. Peptide solutions were added to a 50% human serum solution (1:1 ratio, final serum concentration is 25%) and incubated at 37 ˚C for 0, 6. 24, 30 and 48 hours. At 0 hour and selected, variable time points, 20 μL of the mixture was transferred to a new tube and 40 μL 96% ethanol added to quench the reaction. The quenched aliquots were cooled at 4 ˚C for 30min and spun down at 14000 RPM for 5 min. The supernatant (around 60 μL) was collected and could be frozen at -80 ˚C until ready for analysis. Immediately after thawing, quenched samples were run on TOF-HPLC on a C18 column to assess peptide degradation.

**Viability assay.** Cells were passaged and resuspended in 10% FBS DMEM with suggested antibiotics at 4000 cells per well in 96-well plates. Cells were incubated at 37 °C in 5% carbon dioxide overnight. The medium was changed to 1% FBS DMEM and peptides of different concentrations or 1% DMSO were added to each well. Cells were incubated at 37 °C in 5% carbon dioxide for 7 days with old media removed and replaced with fresh media and peptides every 24 hours. At the 168-hour point, cells were aspirated and incubated with 110 μL of 10:1 1% FBS in DMEM: MTT solution for 4 hours at 37 °C in 5% carbon dioxide. 100 μL of the SDS-HCl solution was added to each well to dissolve the purple formazan. Plates were incubated overnight, and the absorbance of each well was read at 570 nm wavelength.

**RT-qPCR assay.** Cells were passaged, resuspended in 10% FBS DMEM at 50,000 cells/mL, and the suspension was added (1000 μL) to clear 6-well plates. After incubating for 24 hours at 37 °C in 5% carbon dioxide, 10 μL 100X peptide in DMSO or DMSO in 1% FBS DMEM (990 μL) were added to each well. Cells were incubated for 4 hours at 37 °C in 5% carbon dioxide. Cell suspensions were centrifuged, aspirated, and the mRNA extracted. Extracted mRNA was converted to cDNA before being subjected to qPCR on an Applied Biosystems StepOnePlus Real Time PCR System. The p value was calculated by an two-tailed unpaired t test.[13] The primers used in qPCR can be found in Table S1. All data points are derived from technical triplicates and biological duplicates.

| Peptide | Sequence (N'-C') |
|---|---|
| CREBLL-tide | Ac-DCGNILPSDIMDFVLKNTPY-$NH_2$<br>(AEEAc)TDSQKRREILSRRPS(phos)YRKILNDLSSDAPG-$NH_2$ |
| CREBLL-tide-Neg | Ac-DCGNILPSDIMDAVLKNTPY-$NH_2$<br>TDSQKRREILSRRPSYRKILNDLSSDAPG-$NH_2$ |
| CREBLL-tide (no linker) | Ac-DCGNILPSDIMDFVLKNTPY-$NH_2$<br>TDSQKRREILSRRPS(phos)YRKILNDLSSDAPG-$NH_2$ |
| CREBLL-tide (short linker) | Ac-DCGNILPSDIMDFVLKNTPY-$NH_2$<br>(βAla)TDSQKRREILSRRPS(phos)YRKILNDLSSDAPG-$NH_2$ |
| CREBLL-tide (medium linker) | Ac-DCGNILPSDIMDFVLKNTPY-$NH_2$<br>(6-Ahx)TDSQKRREILSRRPS(phos)YRKILNDLSSDAPG-$NH_2$ |
| CLIP6-CREBLL-tide | Ac-KVRVRVRV$^{D}$PPTRVRERVKGGDCGNILPSDIMDFVLKNTPY-$NH_2$<br>(AEEAc)TDSQKRREILSRRP-S(phos)YRKILNDLSSDAPG-$NH_2$ |
| CLIP6-CREBLL-Neg | Ac-KVRVRVRV$^{D}$PPTRVRERVKGGDCGNILPSDIMDAVLKNTPY-$NH_2$<br>TDSQKRREILSRRPSYRKILNDLSSDAPG-$NH_2$ |
| AEEAc-pKID$_{119-147}$ | Ac-AEEAc-TDSQKRREILSRRPS(phos)YRKILNDLSSDAPG-$NH_2$ |
| MLL$_{2840-2858}$ | Ac- DCGNILPSDIMDFVLKNTPY-$NH_2$ |

| | |
|---|---|
| Biotin-CREBLL-tide | Biotin-(AEEAc)DCGNILPSDIMDFVLKNTPY-$NH_2$<br>(AEEAc)TDSQKRREILSRRPS(phos)YRKILNDLSSDAPG-$NH_2$ |

| | |
|---|---|
| Biotin-CREBLL-NEG | Biotin-(AEEAc)DCGNILPSDIMDAVLKNTPY-$NH_2$<br>TDSQKRREILSRRPSYRKILNDLSSDAPG-$NH_2$ |
| Biotin-AEEAc-MLL | Biotin-AEEAc-DCGNILPSDIMDFVLKNTPY-$NH_2$ |
| Biotin-(β-Ala)-MLL | Biotin- (β-Ala) -DCGNILPSDIMDFVLKNTPY-$NH_2$ |
| FITC-CREBLL | FITC- (β-Ala)DCGNILPSDIMDFVLKNTPY-$NH_2$<br>(AEEAc)TDSQKRREILSRRPS(phos)YRKILNDLSSDAPG-$NH_2$ |
| FITC-CREBLL-tide-Nophos | FITC- (β-Ala)DCGNILPSDIMDFVLKNTPY-$NH_2$<br>(AEEAc)TDSQKRREILSRRPSYRKILNDLSSDAPG-$NH_2$ |
| FITC-CREBLL-tide-F2852A | FITC- (β-Ala)DCGNILPSDIMDAVLKNTPY-$NH_2$<br>(AEEAc)TDSQKRREILSRRPS(phos)YRKILNDLSSDAPG-$NH_2$ |
| FITC-CREBLL-tide-Neg | FITC- (β-Ala)DCGNILPSDIMDAVLKNTPY-$NH_2$<br>TDSQKRREILSRRPSYRKILNDLSSDAPG-$NH_2$ |
| FITC-CLIP6-CREBLL-tide | FITC-(βAla)-KVRVRVRV$^{D}$PPTRVRERVKGGDCGNILPSDIMDFVLKNTPY-$NH_2$<br>(AEEAc)TDSQKRREILSRRPS(phos)YRKILNDLSSDAPG-$NH_2$ |
| FITC-pKID$_{119\text{-}147}$ | FITC-(AEEAc)-TDSQKRREILSRRPS(phos)YRKILNDLSSDAPG-$NH_2$ |
| FITC-MLL$_{2840\text{-}2858}$ | FITC-(β-Ala) DCGNILPSDIMDFVLKNTPY-$NH_2$ |

| | |
|---|---|
| FITC-TAT-CREBLL-tide | FITC- (βAla)-<br>GRKKRRQRRRGGDCGNILPSDIMDFVLKNTPY-$NH_2$<br>(AEEAc)TDSQKRREILSRRPS(phos)YRKILNDLSSDAPG-$NH_2$ |

**Table S1.** The sequences of the peptides used in this study. The differences between CREBLL-tide and CREBLL-Neg mutants are highlighted in red.

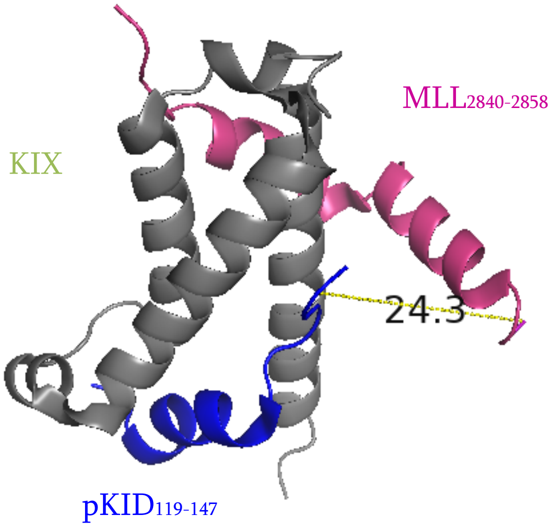


**Figure S2. The NMR structure of MLL-KIX-pKID (2LXT).** The yellow line represents the distance between the Cα atoms of the T119 pKID residue and A2841 MLL residue. This is the structure of pose 5 in the Table S2.

| 2LXT Chain | Distance (Å) |
|---|---|
| 1 | 13.6 |
| 2 | 19.5 |
| 3 | 16.4 |
| 4 | 19.1 |
| 5 | 21.8 |
| 6 | 19.8 |
| 7 | 22.5 |
| 8 | 17 |
| 9 | 25.4 |
| 10 | 18.3 |
| 11 | 15.1 |
| 12 | 17.5 |
| 13 | 23.2 |
| 14 | 25.4 |
| 15 | 23.5 |
| 16 | 19.6 |
| 17 | 16.8 |
| 18 | 13.6 |
| 19 | 21.6 |
| 20 | 24.1 |
| Average | 19.6 |

| | |
|---|---|
| SD | 3.7 |
| Min | 13.6 |
| Max | 25.4 |

**Table S2**. Distance between the Cα atoms of the T119 pKID residue and A2841 MLL residue from NMR structure (PDB: 2LXT). The different distances are from 20 different poses of NMR structure.

| Name | Structure | Length (Å) |
|---|---|---|
| Short linker: β-Alanine | 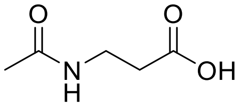 | 9.0-10.0 |
| Medium linker: 6-Ahx | 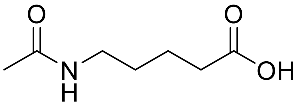 | 11.6-12.6 |
| Long linker: AEEAc | 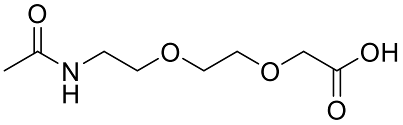 | 15.8-16.8 |

**Table S3.** The length and structure of different linkers for the CREBLL-tides. The distance was measured from the N atom on T119 of pKID to the S atom on C2841 of $MLL_{2840-2858}$ after constructing the linkers from the N-terminal amine of pKID to the cysteine side chain of $MLL_{2840-2858}$.

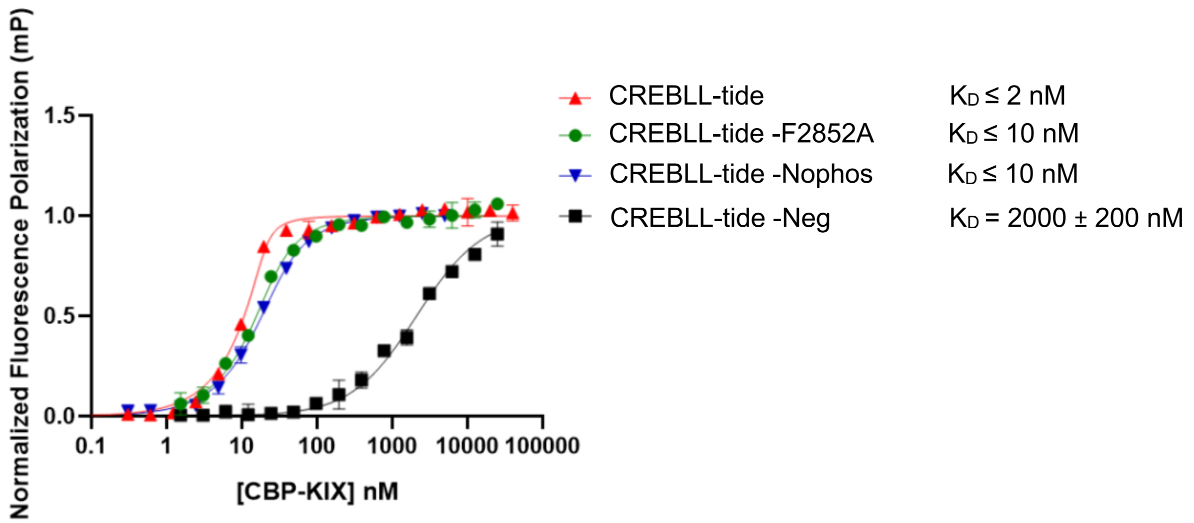


**Figure S3**. The binding affinity of CREBLL-tide mutants to CBP KIX. Error is SD of independent experimental triplicates. See Table S1 or Figure S14-S31 for structures.

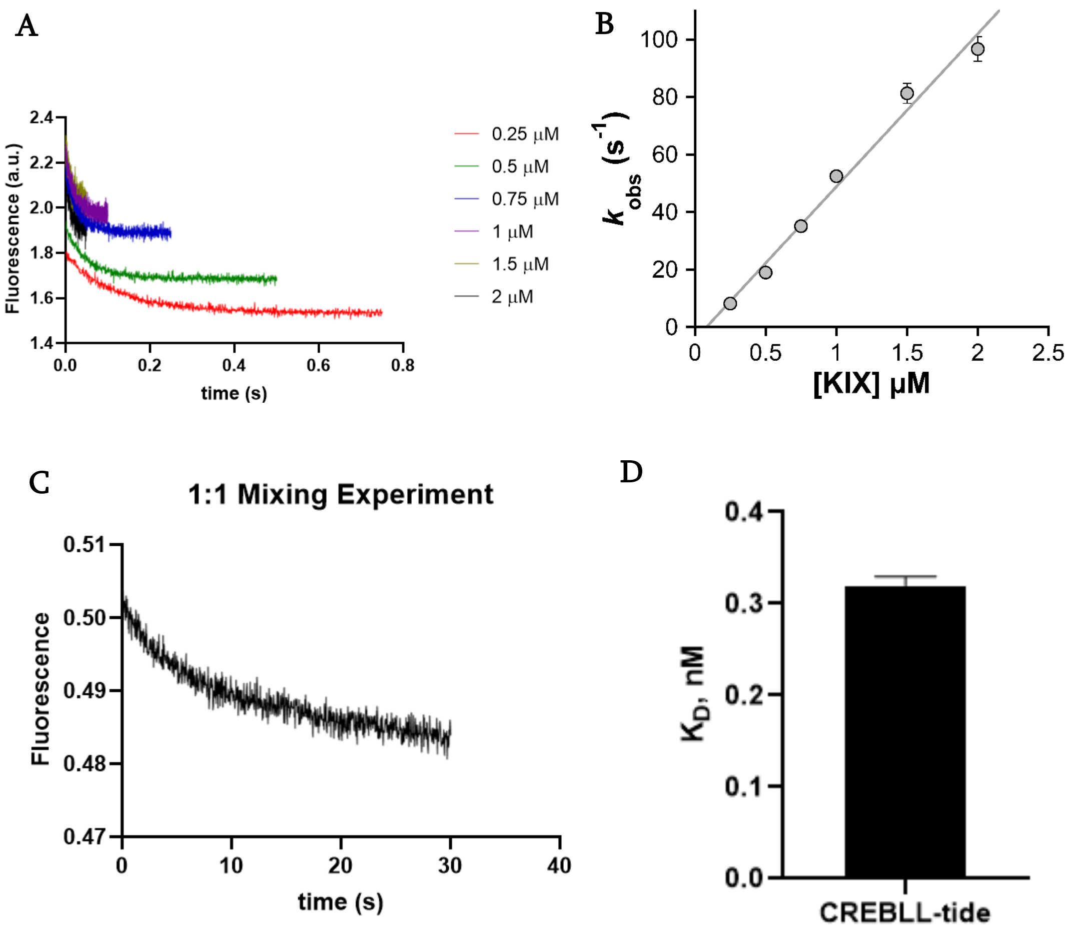


**Figure S4.** Kinetic data of CREBLL-tide binding to KIX as measured by stopped-flow fluorescence. (A) The stopped-flow data of increasing concentration of KIX with 25 nM FITC-CREBLL. Similar experiments were performed with different concentrations to determine kon. (B) The kobs data gained from concentration dependence experiments, were plotted with KIX concentration [KIX] and fit a linear equation to determine the value of kon. Error bars are standard deviations derived from three experimental replicates. (C) The 1:1 mixing experiment of 5 nM FITC-CRBLL-tide with 5 nM CBP-KIX. The data were fit to calculate the koff. All data calculation was average of 10-15 1:1 association experiment. (D) The $K_D$ of CREBLL-tide for CBP-KIX which was calculated based on the $k_{on}$ and $k_{off}$ (KD = $k_{off}/k_{on}$). Error represents SD

| Peptide | $k_{on}$ (μM-1•s-1) | $k_{off}$ (s-1) | $K_D$ (nM) |
|---|---|---|---|
| FITC-CREBLL-tide | 53±3 | 0.017±0.002 | 0.32±0.01 |

**Table S4.** The kon, koff, and $K_D$ were calculated from the data of stopped-flow experiments.

$K_D$=$k_{off}$/$k_{on}$. Mean ± SD. Error is SD from 10-15 traces of experimental triplicates.

A

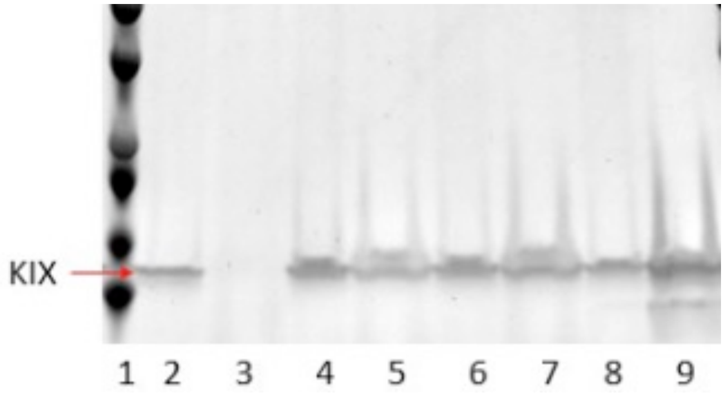


1. Ladder
2. FT of blank
3. PD of blank
4. FT of Biotin-(β-Ala)-MLL
5. PD of Biotin-(β-Ala)-MLL
6. FT of Biotin-AEEAc-MLL
7. PD of Biotin-AEEAc-MLL
8. FT of Biotin-CREBLL-tide
9. PD of Biotin-CREBLL-tide

B

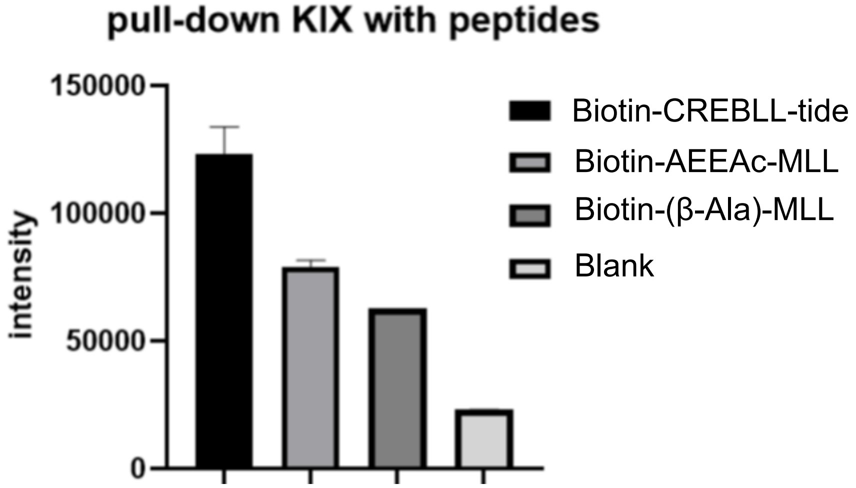


**Figure S5.** Pulldown of KIX with biotin-labeled peptides. (A) The SDS-PAGE gel after Coomassie blue staining. FT represents the flow through of each sample without beads. PD represents the Pull-down sample with beads. The red arrow represents where the KIX is. (B) The quantification of the SDS-PAGE of each PD sample analyzed by Image J. Error is SD from different exposure time for the blots.

CBP

| Contrast | LogFC | p-value |
|---|---|---|
| SUM149.WT vs. MCF10A.WT | 0.010 | 0.997 |
| SUM190.WT vs. MCF10A.WT | ▴0.950 | 9.669e-6 |
| VARI068.WT vs. MCF10A.WT | ▴1.309 | 1.000e-6 |
| MDA231.WT vs. MCF10A.WT | 0.181 | 0.490 |
| MCF7.WT vs. MCF10A.WT | ▴1.430 | 1.000e-6 |

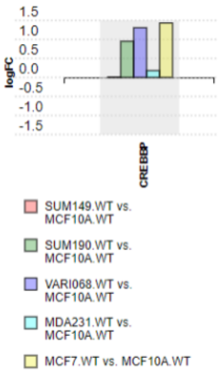


p300

| Contrast | LogFC | p-value |
|---|---|---|
| SUM149.WT vs. MCF10A.WT | ▴1.657 | 1.000e-6 |
| SUM190.WT vs. MCF10A.WT | ▴1.598 | 1.000e-6 |
| VARI068.WT vs. MCF10A.WT | ▴2.438 | 1.000e-6 |
| MDA231.WT vs. MCF10A.WT | ▴1.333 | 1.000e-6 |
| MCF7.WT vs. MCF10A.WT | ▴1.684 | 1.000e-6 |

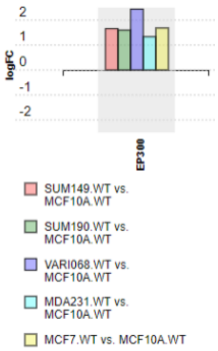


**Figure S6.** A comparison of CBP and p300 mRNA levels in various breast cancer cell line obtained from RNA-seq analysis. LogFC (log2 fold change) represents the effect size estimate. The red number represents the overexpression of the gene in that specific cell line.

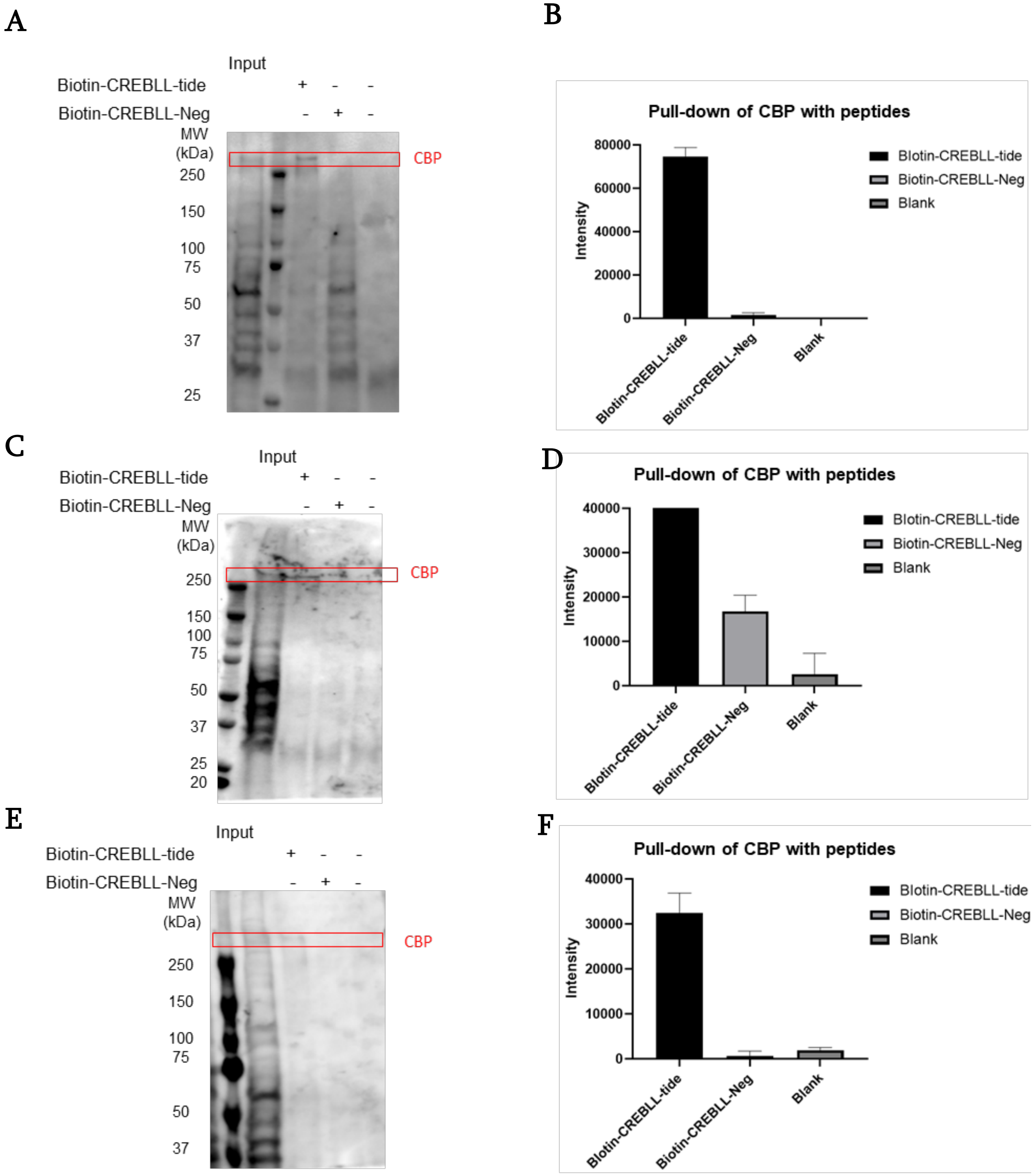
A
Input
Biotin-CREBLL-tide + - -
Biotin-CREBLL-Neg - + -
MW (kDa)
250
150
100
75
50
37
25
CBP
B
Pull-down of CBP with peptides
Intensity
80000
60000
40000
20000
0
Biotin-CREBLL-tide
Biotin-CREBLL-Neg
Blank
C
Input
Biotin-CREBLL-tide + - -
Biotin-CREBLL-Neg - + -
MW (kDa)
250
150
100
75
50
37
25
20
CBP
D
Pull-down of CBP with peptides
Intensity
40000
30000
20000
10000
0
Biotin-CREBLL-tide
Biotin-CREBLL-Neg
Blank
E
Input
Biotin-CREBLL-tide + - -
Biotin-CREBLL-Neg - + -
MW (kDa)
250
150
100
75
50
37
CBP
F
Pull-down of CBP with peptides
Intensity
40000
30000
20000
10000
0
Biotin-CREBLL-tide
Biotin-CREBLL-Neg
Blank

**Figure S7.** (A) Western blot for endogenous, full-length CBP from VARI068 whole cell lysates showing affinity pulldown of CBP via biotinylated CREBLL-tide but not Biotin-CREBLL-Neg. 10 μM Biotin-CREBLL-tide and Biotin-CREBLL-Neg were bound to the resin and incubated with the cell lysate for 1 hour. (B) The quantification of the intensity of CBP pulled down with biotinylated peptides on Western Blot by Image J. Error is SD from different exposure time for the bolts. (C), (D), (E) and (F) are the data of other replicates.

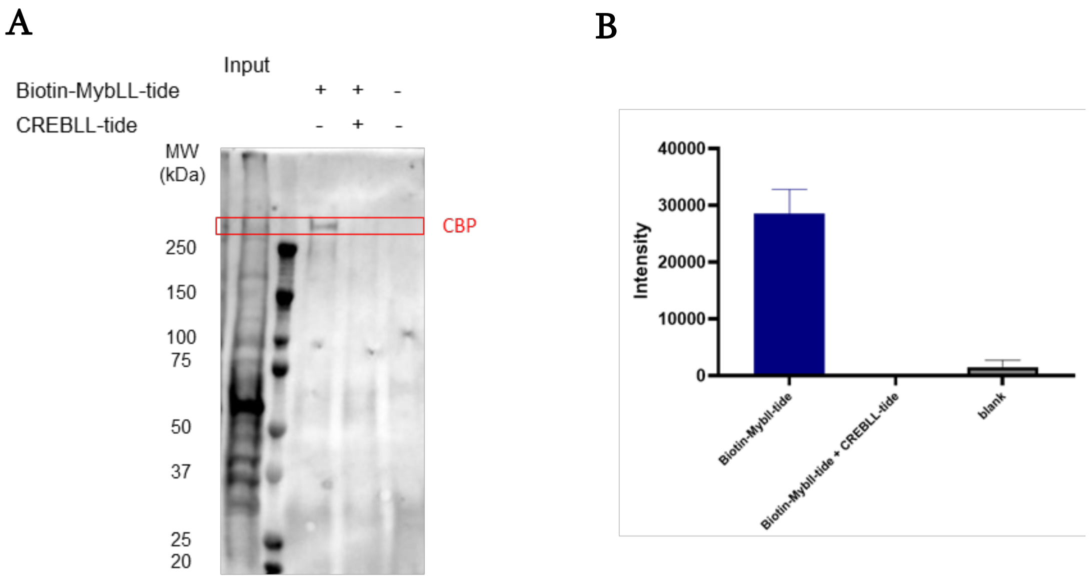


**Figure S8.** (A) Western blot for endogenous, full-length CBP from VARI068 whole cell lysates showing affinity pulldown of CBP via biotinylated MybLL-tide and competitive inhibition of Biotin-MybLL-tide by CREBLL-tide. 10 μM Biotin-MybLL-tide with or without 50 μM of CREBLL-tide were bound to the resin and incubated with the cell lysate for 1 hour. (B) The quantification of the intensity of CBP pulled down with biotinylated peptides on Western Blot by Image J. Error is SD from different exposure time for the bolts.

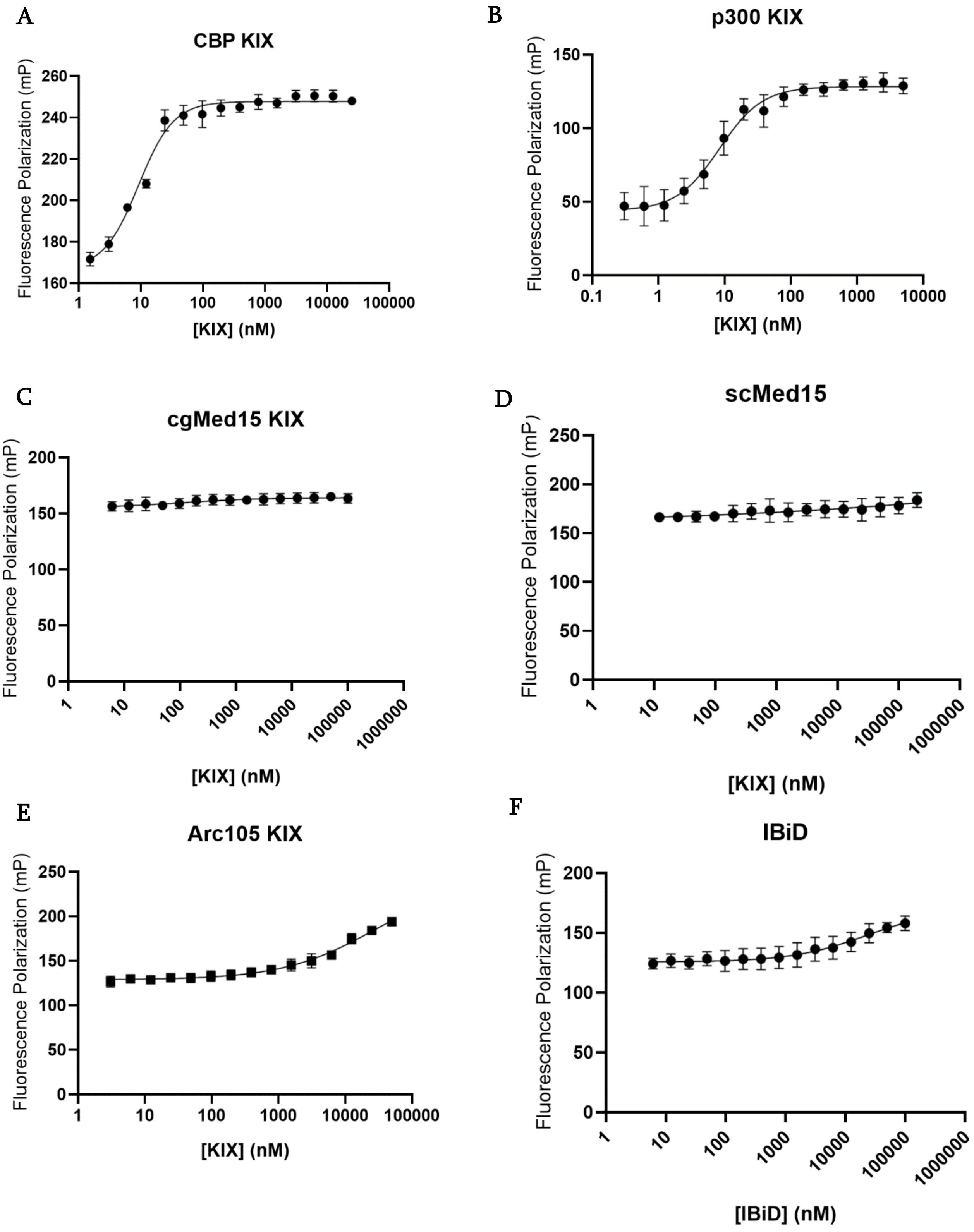
A
CBP KIX
Fluorescence Polarization (mP)
260
240
220
200
180
160
1
10
100
1000
10000
100000
[KIX] (nM)
B
p300 KIX
Fluorescence Polarization (mP)
150
100
50
0
0.1
1
10
100
1000
10000
[KIX] (nM)
C
cgMed15 KIX
Fluorescence Polarization (mP)
200
150
100
50
0
1
10
100
1000
10000
100000
1000000
[KIX] (nM)
D
scMed15
Fluorescence Polarization (mP)
250
200
150
100
50
0
1
10
100
1000
10000
100000
1000000
[KIX] (nM)
E
Arc105 KIX
Fluorescence Polarization (mP)
250
200
150
100
50
0
1
10
100
1000
10000
100000
[KIX] (nM)
F
IBiD
Fluorescence Polarization (mP)
200
150
100
50
0
1
10
100
1000
10000
100000
1000000
[IBiD] (nM)

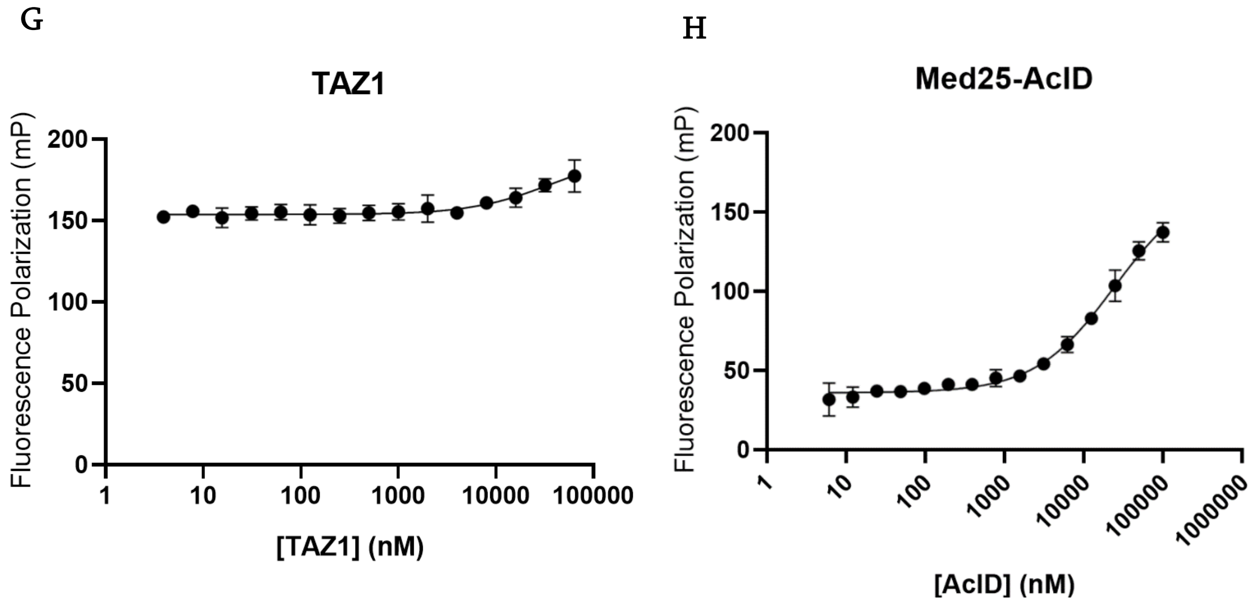


**Figure S9.** Direct-binding fluorescence polarization data for CREBLL-tide with the indicated activator-binding domains. 20 nM FITC-CREBLL-tide binding to (A) CBP KIX, (B) p300 KIX, (C) cgMED15 KIX, (D) scMED15 KIX, (E) Arc105 KIX, (F) CBP IBiD, (G) p300 TAZ1, (H) MED25 AcID. Error bars are SD calculated from technical triplicates of independent experimental triplicates.

| Protein domain | $K_D$ of FITC-CREBLL-tide (nM) |
|---|---|
| CBP KIX | 0.32±0.04* |
| Arc105 KIX | 8800±500 |
| scMED15 KIX | >200,000 |
| cgMED15 KIX | >200,000 |
| IBiD | >100,000 |
| TAZ1 | >100,000 |
| MED25 AcID | 19000±4000 |

**Table S5.** KD for FITC-CREBLL-tide, and FITC-MybLL-tide with various domains. The data of CREBLL-tide is derived from above direct-binding fluorescence polarization experiments.

* denotes data for CBP interaction derived from above stopped-flow fluorescence experiments. Mean ± SD. Error is SD from independent experimental triplicates.

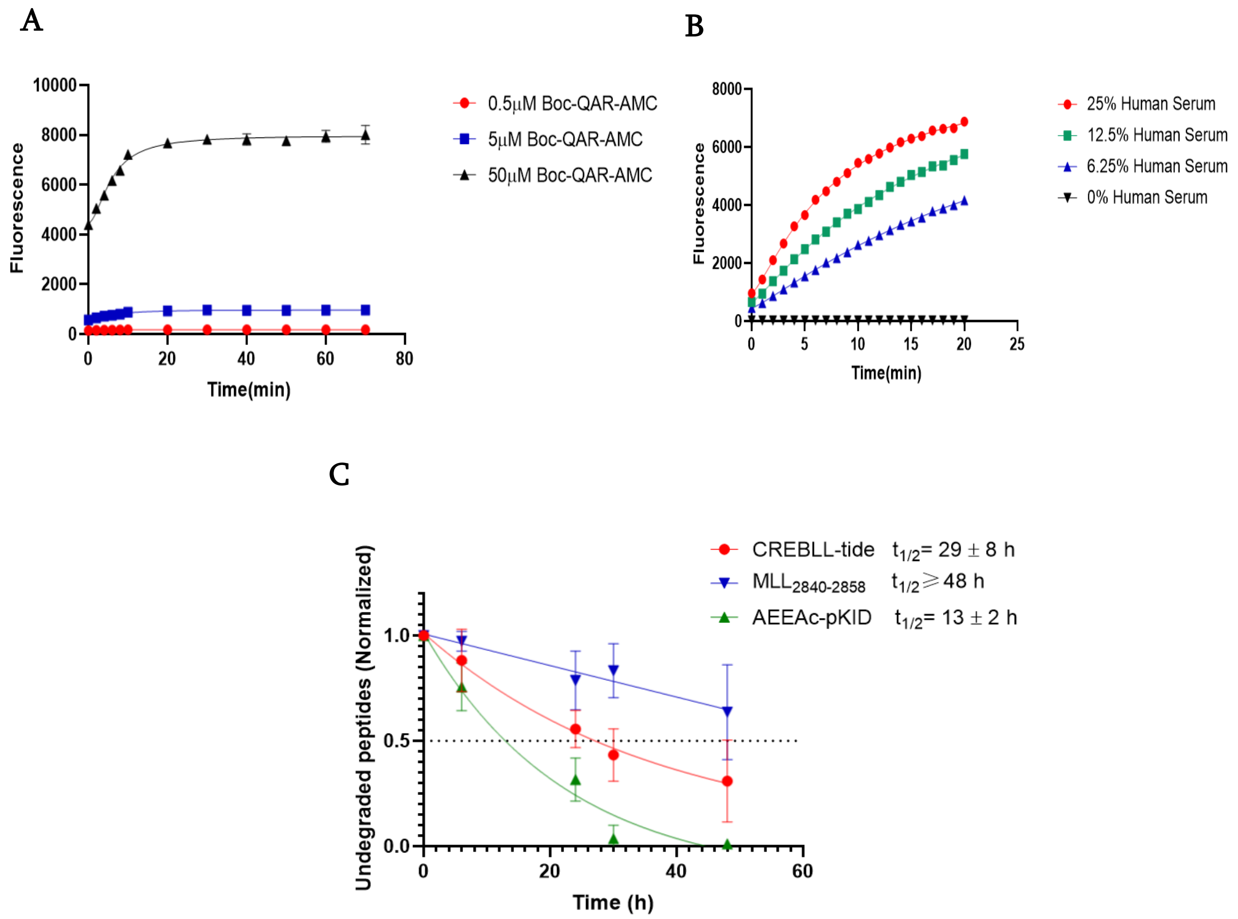


**Figure S10**. The stability of CREBLL-tides in human serum. (A) The fluorescence change of Boc-QAR-AMC with different concentrations at different time points in 25% human serum. Experiments were performed in technical triplicates. Error is the standard deviation of technical triplicates. (B) The fluorescence change of 50 μM Boc-QAR-AMC at different time points in human serum with increasing concentration. Experiments were performed in technical triplicates. Error is the standard deviation of technical triplicates. (C) The undegraded peptides after incubating with 25% human serum for different times. Error is SD

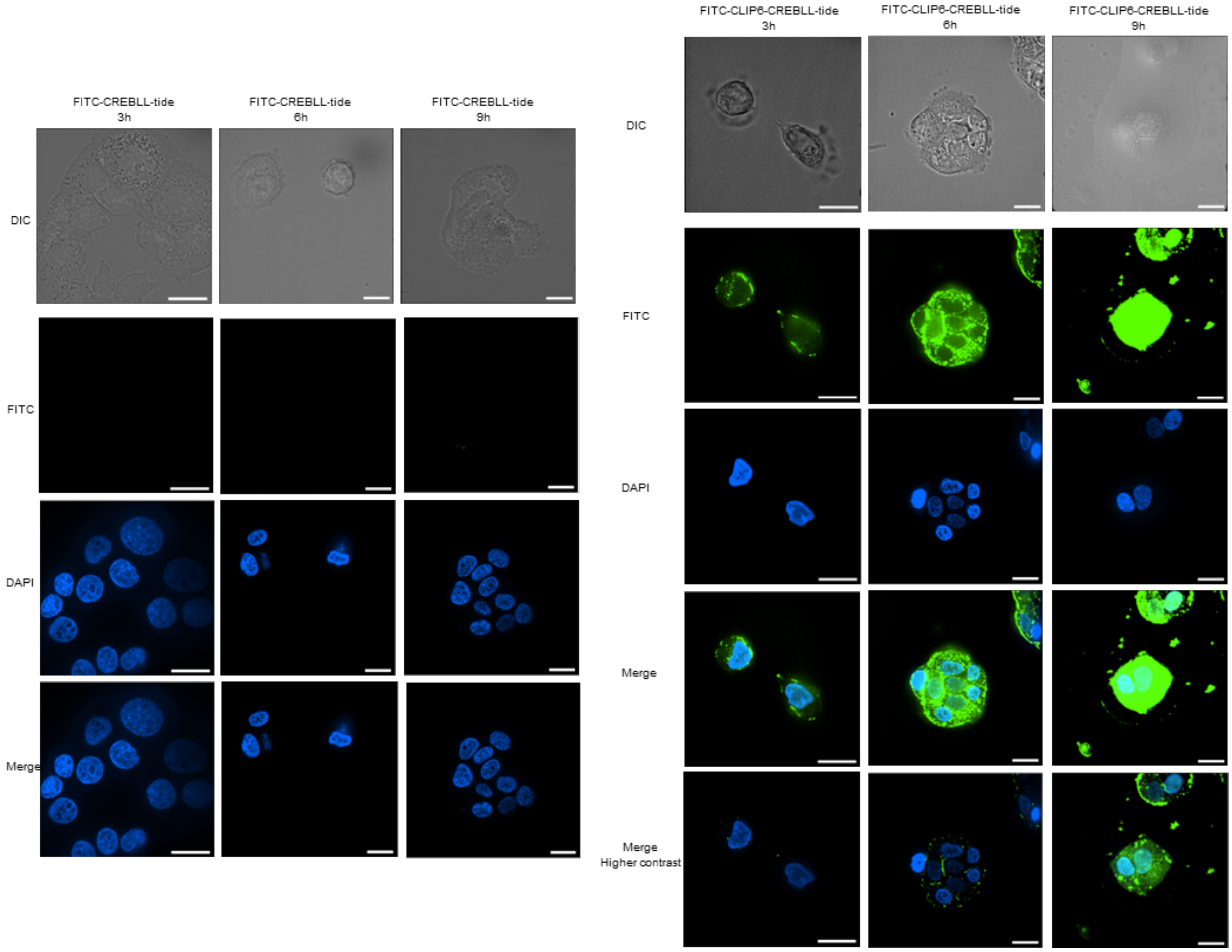


**Figure S11.** The DIC, FITC and DAPI channel confocal images of VARI068 cells incubating with 10 μM FITC-CREBLL-tide (left) and FITC-CLIP6-CREBLL-tide (right) for 3 h, 6h and 9h. The Merge images are merged by DAPI and FITC channel. Merged images with higher contrast of FITC-CLIP6-CREBLL-tide were also showed. Blue staining in the DAPI and Merge images represent the size and location of the nucleus. Green dots show the location of the FITC labelled peptides in cells. Each scale bar represents 20-micron length in the image.

A

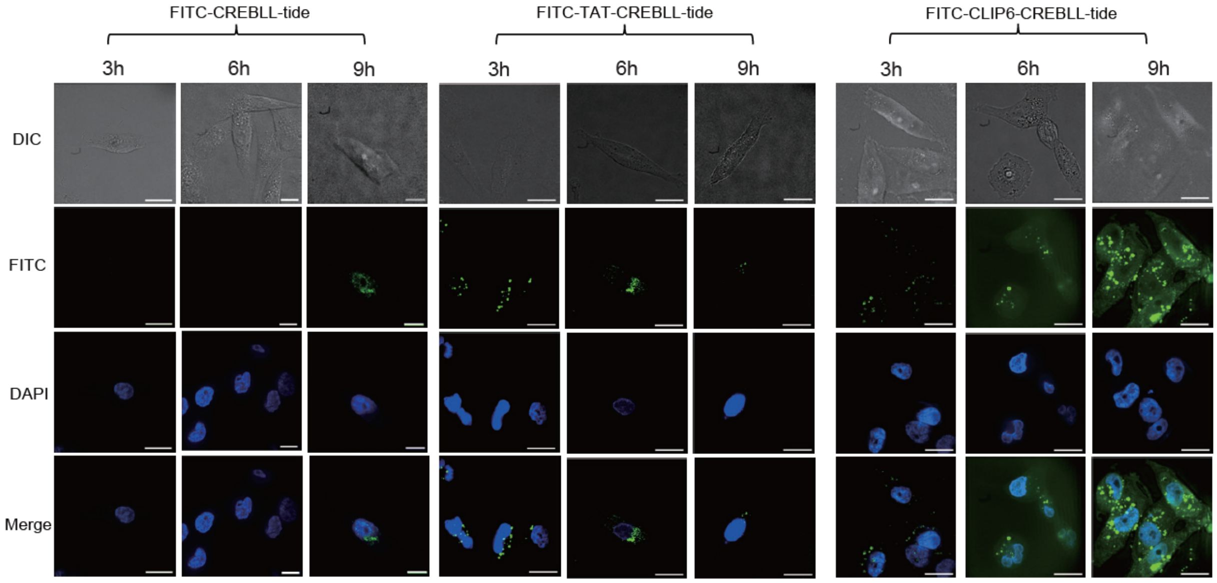

FITC-CREBLL-tide
FITC-TAT-CREBLL-tide
FITC-CLIP6-CREBLL-tide
3h
6h
9h
3h
6h
9h
3h
6h
9h
DIC
FITC
DAPI
Merge


B

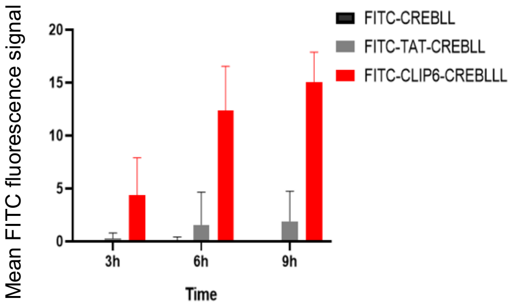

Mean FITC fluorescence signal
20
15
10
5
0
3h
6h
9h
Time
FITC-CREBLL
FITC-TAT-CREBLL
FITC-CLIP6-CREBLLL

**Figure S12**. The cell permeability of CREBLL-tides in MDA-MB-231cells. (A) The DIC, FITC and DAPI channel confocal images of MDA-MB-231 cells incubating with 10 μM FITC-CREBLL-tide (left), FITC-TAT-CREBLL-tide (middle), and FITC-CLIP6-CREBLL-tide (right) for 3 h, 6h and 9h. The Merge images are merged by DAPI and FITC channelBlue staining in the DAPI and Merge images represent the size and location of the nucleus. Green dots show the location of the FITC labelled peptides in cells. Each scale bar represents 20-micron length in the image. (B)The quantification of the mean fluorescence intensity of the peptides in the nucleus of cells at different times. The mean FITC fluorescence signal within the nucleus was taken and the bar graph represents the average of over 10 cells. Error is the standard deviation from average fluorescence signal of biological triplicates.

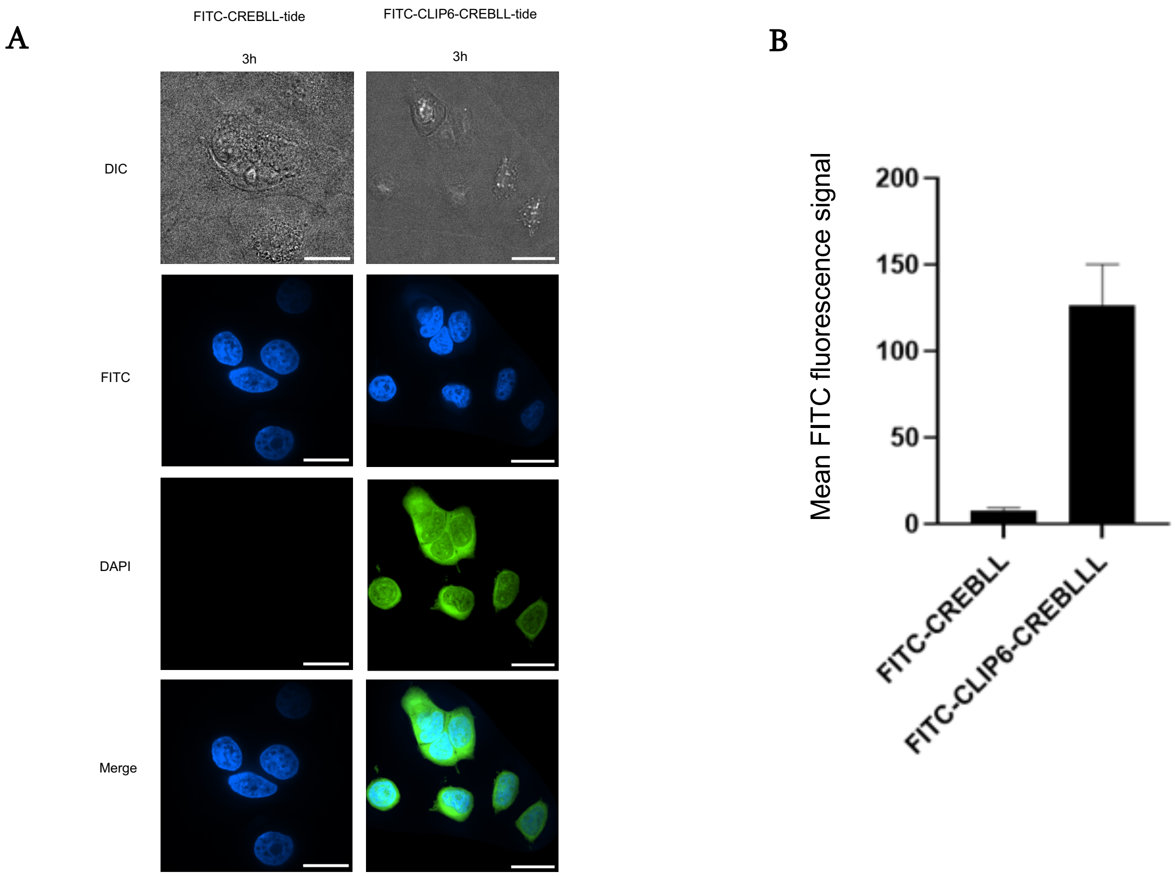


**Figure S13.** The cell permeability of CREBLL-tides in MCF7 cells. (A) The DIC, FITC and DAPI channel confocal images of MCF7 cells incubating with 10 μM FITC-CREBLL-tide (left) and FITC-CLIP6-CREBLL-tide (right) for 3 h. The Merge images are merged by DAPI and FITC channel. Blue staining in the DAPI and Merge images represent the size and location of the nucleus. Green dots show the location of the FITC labelled peptides in cells. Each scale bar represents 20-micron length in the image. (B) The quantification of the mean fluorescence intensity of the peptides in the nucleus of cells at different times. The mean FITC fluorescence signal within the nucleus was taken and the bar graph represents the average of over 10 cells. Error is the standard deviation from average fluorescence signal of biological triplicates.

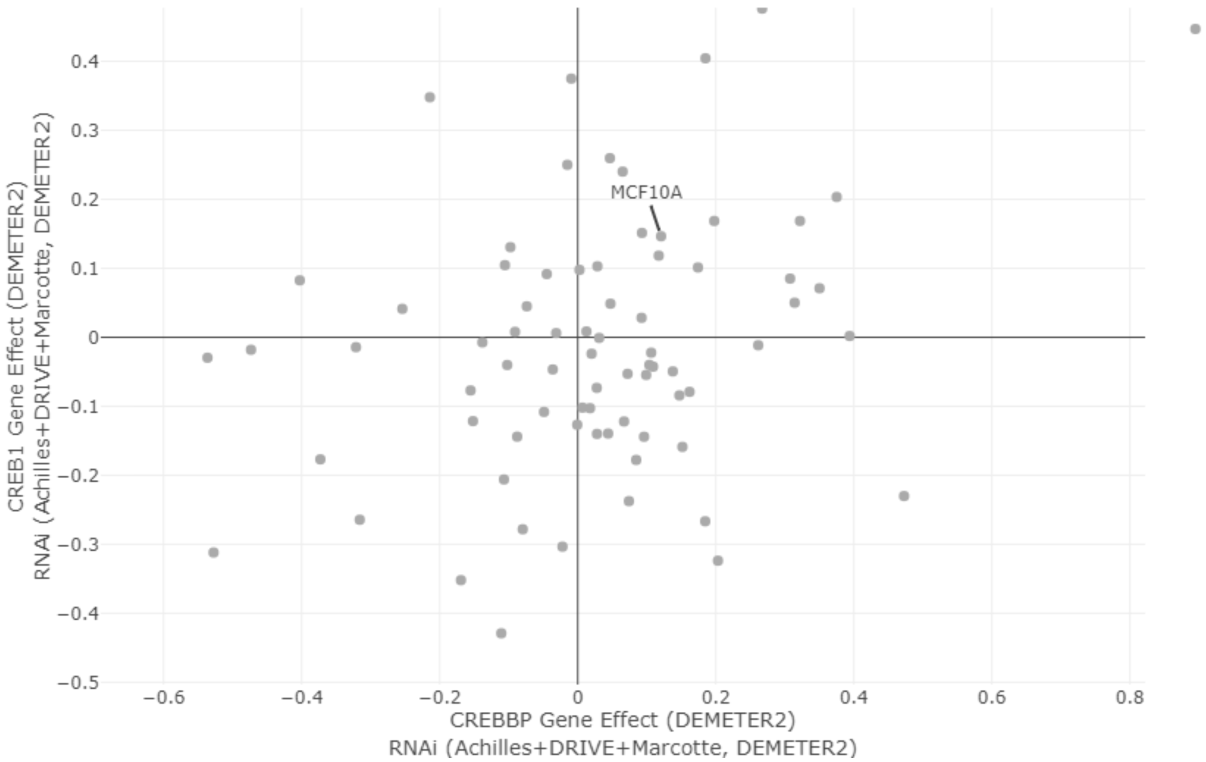


**Figure S14.** The importance of CBP and CREB for different cell lines tested by RNAi assay. The data is adapted from the Depmap website. MCF10A cells are labelled and not dependent on CREB and CBP.[14]

| Primer | Sequence (5’-sequence-3’) |
|---|---|
| RPL19 Forward | ATGTATCACAGCCTGTACCTG |
| RPL19 Reverse | TTCTTGGTCTCTTCCTCCTTG |
| MMP-2 Forward | AGCGAGTGGATGCCGCCTTTAA |
| MMP-2 Reverse | CATTCCAGGCATCTGCGATGAG |
| PTHRP Forward | GAACTCGCTCTGCCTGGTTAGA |
| PTHRP Reverse | GTCCTTGGAAGGTCTCTGCTGA |
| Cyclin D1 Forward | TCTACACCGACAACTCCATCCG |
| Cyclin D1 Reverse | TCTGGCATTTTGGAGAGGAAGTG |
| BCL2 Forward | ATCGCCCTGTGGATGACTGAGT |
| BCL2 Reverse | GCCAGGAGAAATCAAACAGAGGC |

**Table S6.** The sequence of the primers used in the qPCR experiments.

.

| Peptides | Expected MW | Observed MW |
|---|---|---|
| CREBLL-tide | 5919.94 | 5919.93 |
| CREBLL-tide (no linker) | 5774.86 | 5774.87 |
| CREBLL-tide (short linker) | 5845.9 | 5845.91 |
| CREBLL-tide (medium linker) | 5887.95 | 5887.95 |
| CLIP6-CREBLL | 8248.4 | 8248.21 |
| CLIP6-CREBLL-Neg | 7944.31 | 7944.36 |
| AEEAc-pKID | 3624.86 | 3625.88 |
| $MLL_{2840-2858}$ | 2297.08 | 2297.1 |
| Biotin-CREBLL | 6250.08 | 6250.07 |
| Biotin-CREBLL-NEG | 5948.01 | 5948.02 |
| Biotin-AEEAc-MLL | 2625.24 | 2625.23 |
| FITC-CREBLL-tide | 6338 | 6337.98 |
| FITC-CREBLL-tide-Nophos | 6259.02 | 6258.99 |
| FITC-CREBLL-tide-F2852A | 6262.97 | 6262.99 |
| FITC-CREBLL-tide-Neg | 6037.93 | 6037.79 |
| FITC-CLIP6-CREBLL-tide | 8665.43 | 8665.44 |
| FITC-pKID | 3973.29 | 3972.88 |
| FITC-$MLL_{2840-2858}$ | 2714.16 | 2714.01 |
| FITC-TAT-CREBLL-tide | 7829.92 | 7830.89 |

**Table S7.** The expected molecular weight of peptides based on structure, the observed molecular weight was calculated based on the observed molecular weight and ion which was obtained via deconvolution on an Agilent 1260 TOF.

#### Characterization of peptides

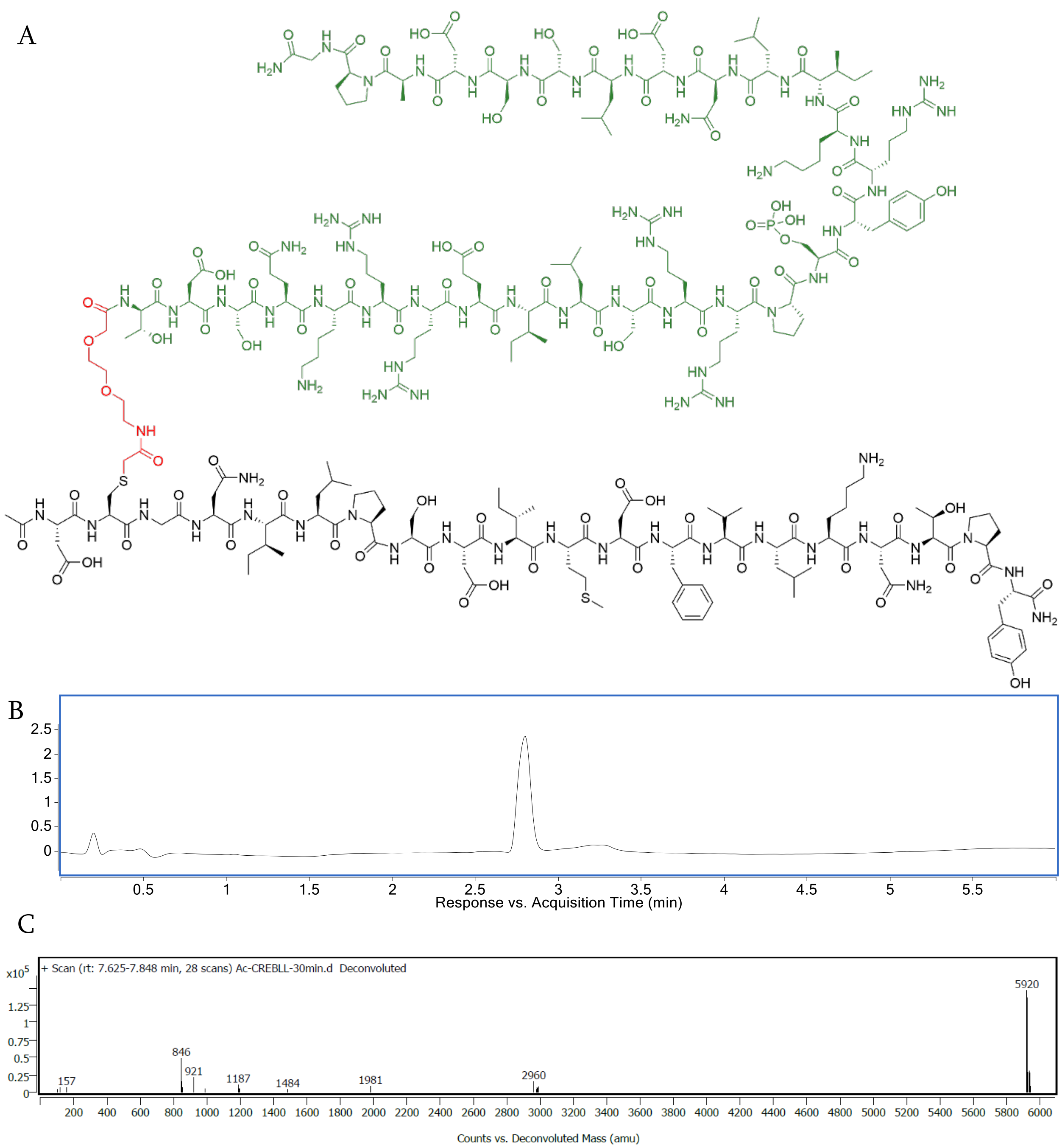


**Figure S15.** Characterization of CREBLL-tide. (A) Chemical structure of the peptide. (B) Analytical trace of CREBLL-tide monitored at 280 nm wavelength. (C) Deconvolution spectra of CREBLL-tide using LC-MS qTOF Mass Spectrometry.

A

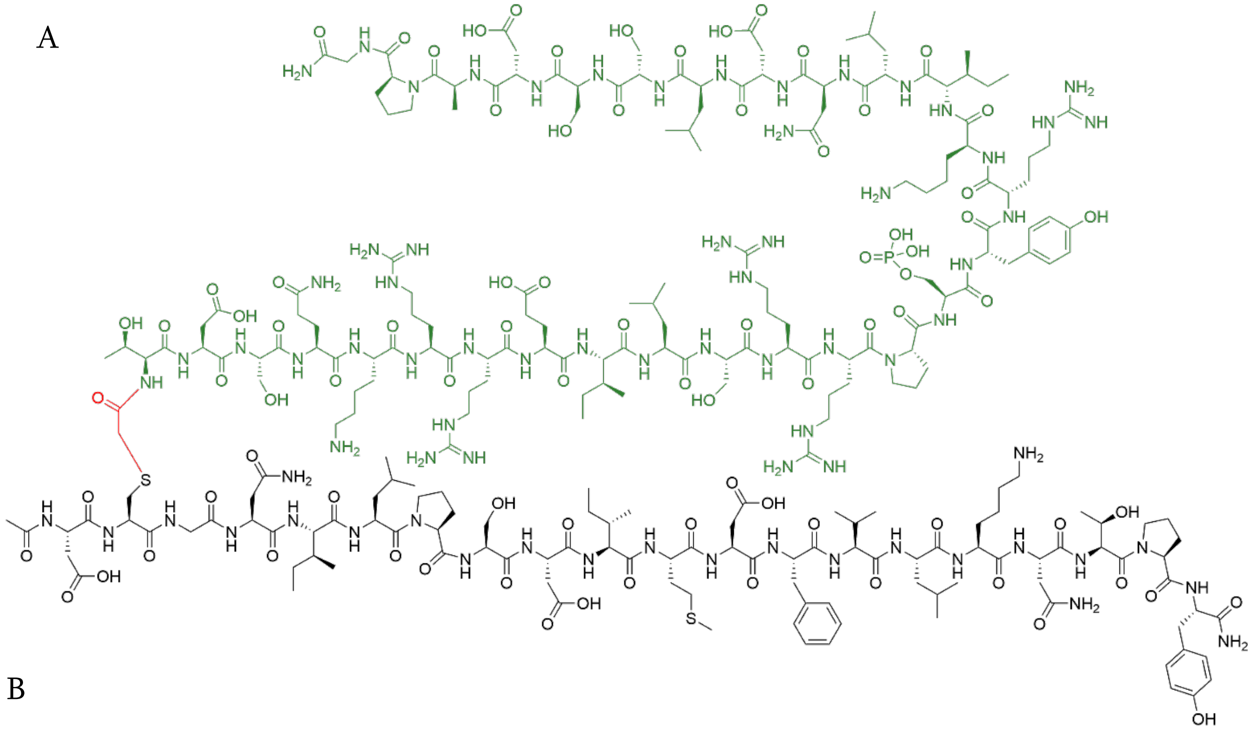

B

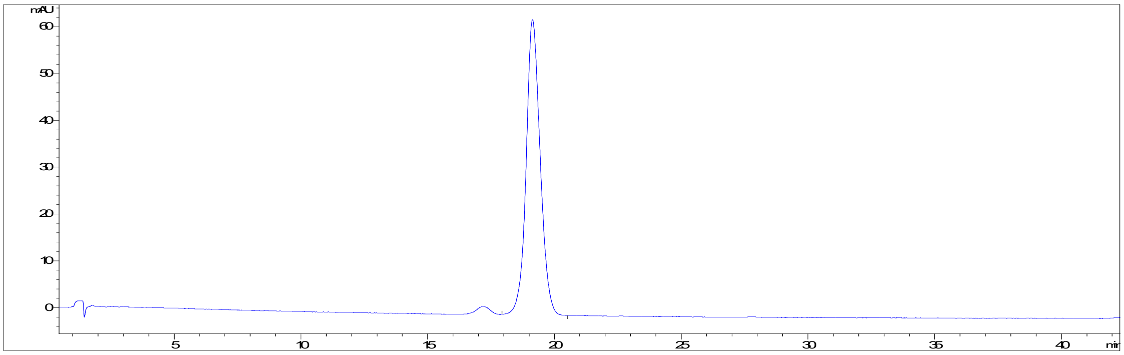
mAU
60
50
40
30
20
10
0
5
10
15
20
25
30
35
40
min

C

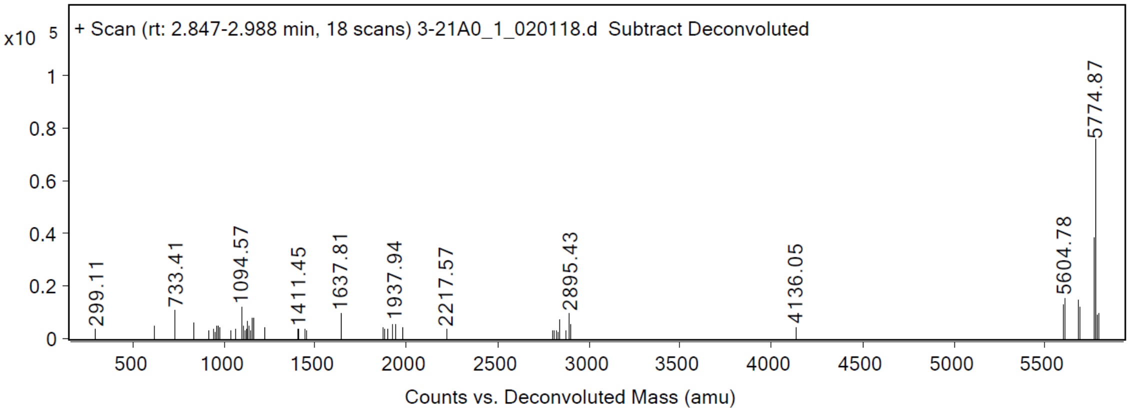
+ Scan (rt: 2.847-2.988 min, 18 scans) 3-21A0_1_020118.d Subtract Deconvoluted
x10 5
1
0.8
0.6
0.4
0.2
0
299.11
733.41
1094.57
1411.45
1637.81
1937.94
2217.57
2895.43
4136.05
5604.78
5774.87
500
1000
1500
2000
2500
3000
3500
4000
4500
5000
5500
Counts vs. Deconvoluted Mass (amu)

**Figure S16.** Characterization of CREBLL-tide (no linker). (A) Chemical structure of the peptide. (B) Analytical trace of CREBLL-tide (no linker) monitored at 280 nm wavelength. (C) Deconvolution spectra of CREBLL-tide (no linker) using LC-MS qTOF Mass Spectrometry.

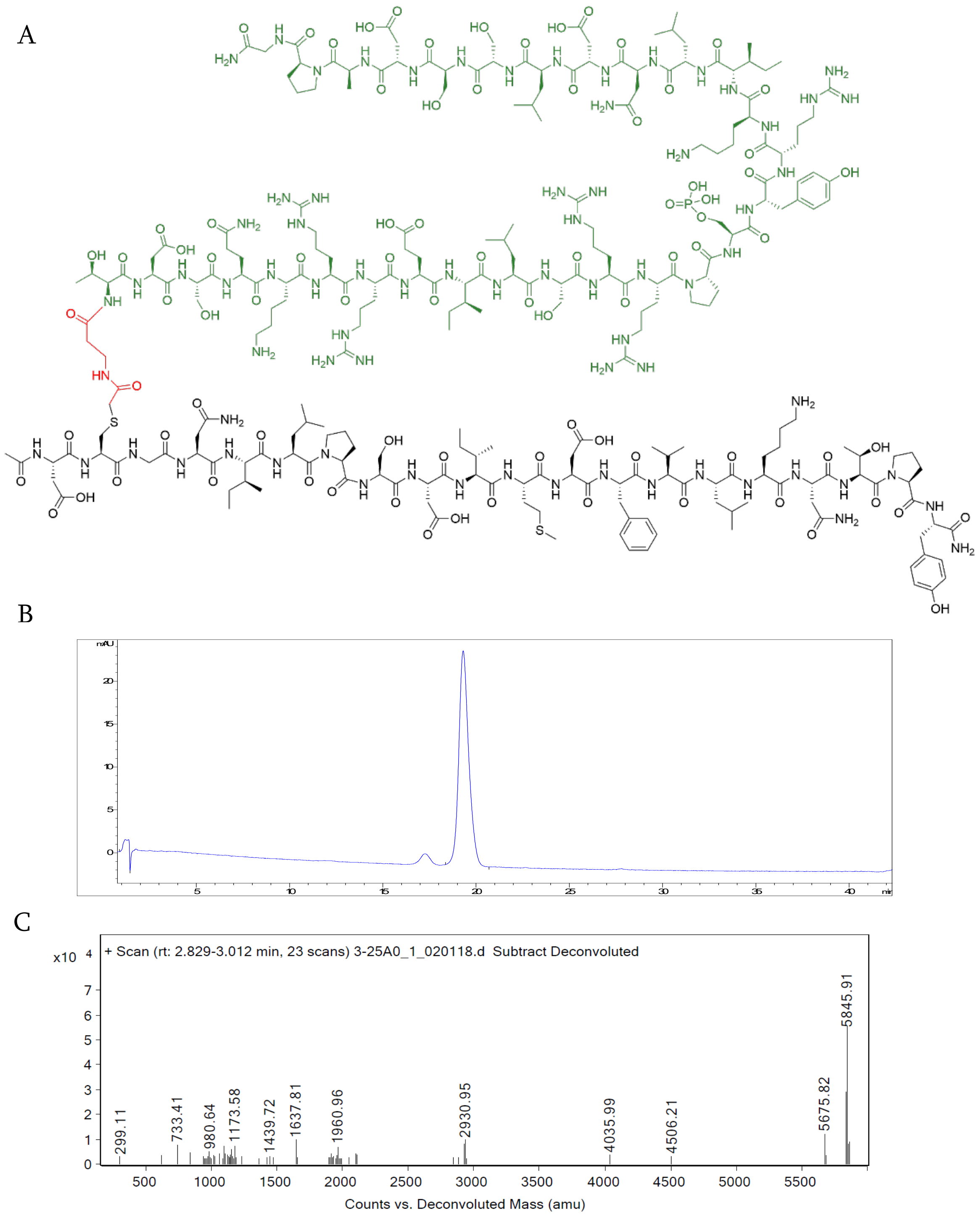


**Figure S17.** Characterization of CREBLL-tide (short linker). (A) Chemical structure of the peptide. (B) Analytical trace of CREBLL-tide (short linker) monitored at 280 nm wavelength. (C) Deconvolution spectra of CREBLL-tide (short linker) using LC-MS qTOF Mass [illegible]

A

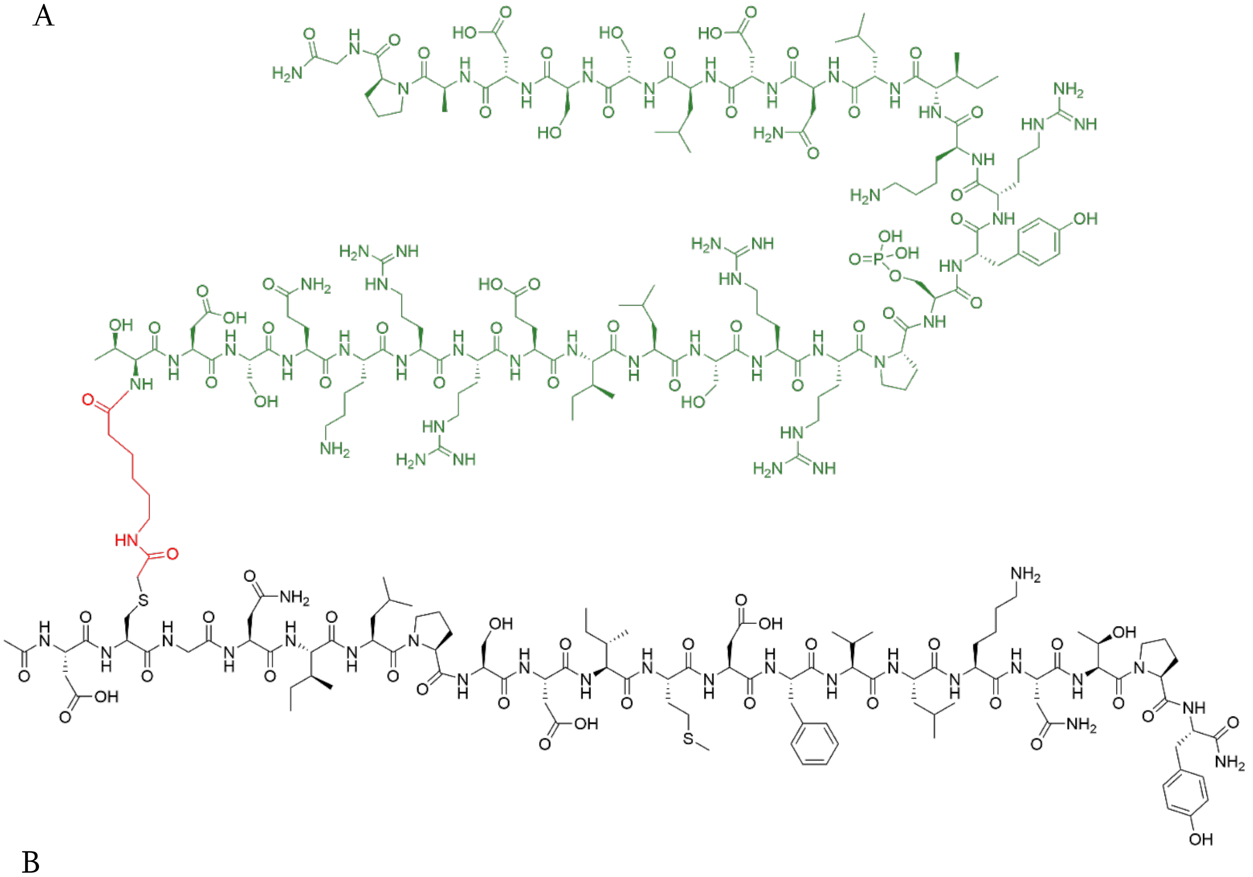

B

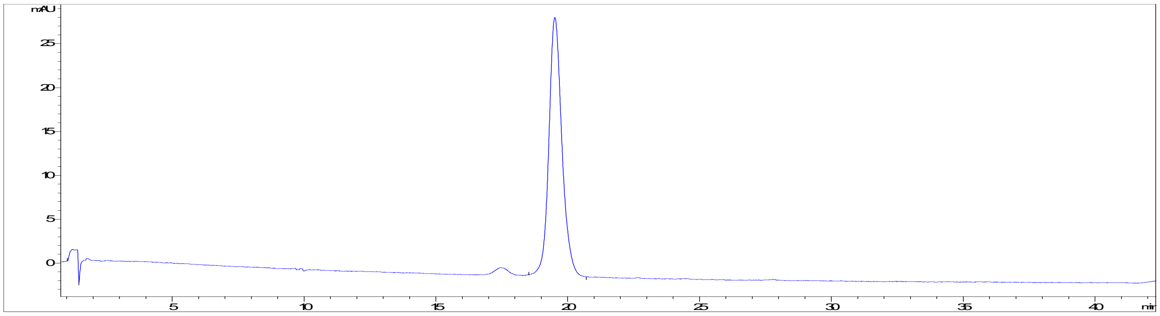

C

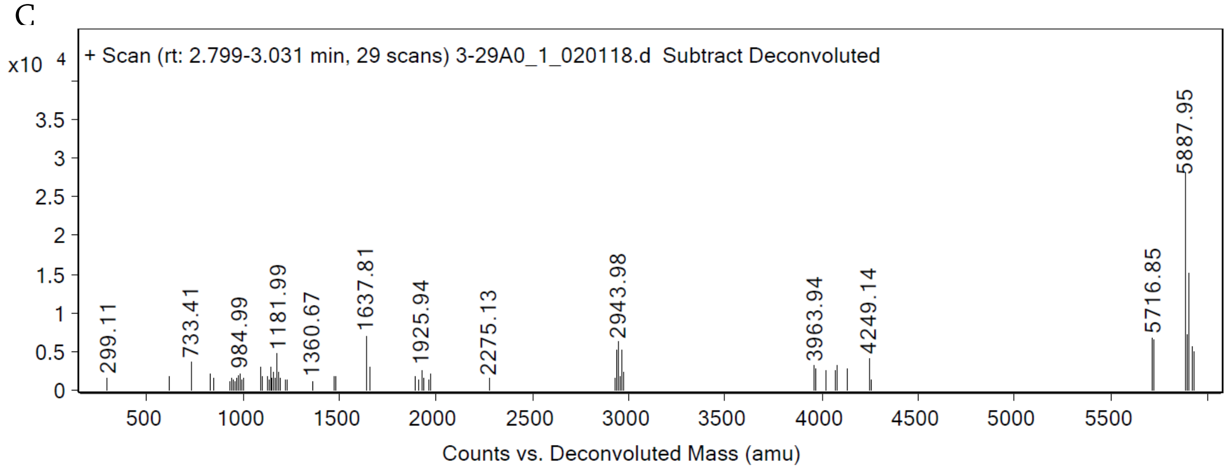
x10 4
+ Scan (rt: 2.799-3.031 min, 29 scans) 3-29A0_1_020118.d Subtract Deconvoluted
299.11
733.41
984.99
1181.99
1360.67
1637.81
1925.94
2275.13
2943.98
3963.94
4249.14
5716.85
5887.95
Counts vs. Deconvoluted Mass (amu)

**Figure S18.** Characterization of CREBLL-tide (medium linker). (A) Chemical structure of the peptide. (B) Analytical trace of CREBLL-tide (medium linker) monitored at 280 nm wavelength. (C) Deconvolution spectra of CREBLL-tide (medium linker) using LC-MS qTOF Mass Spectrometry

A

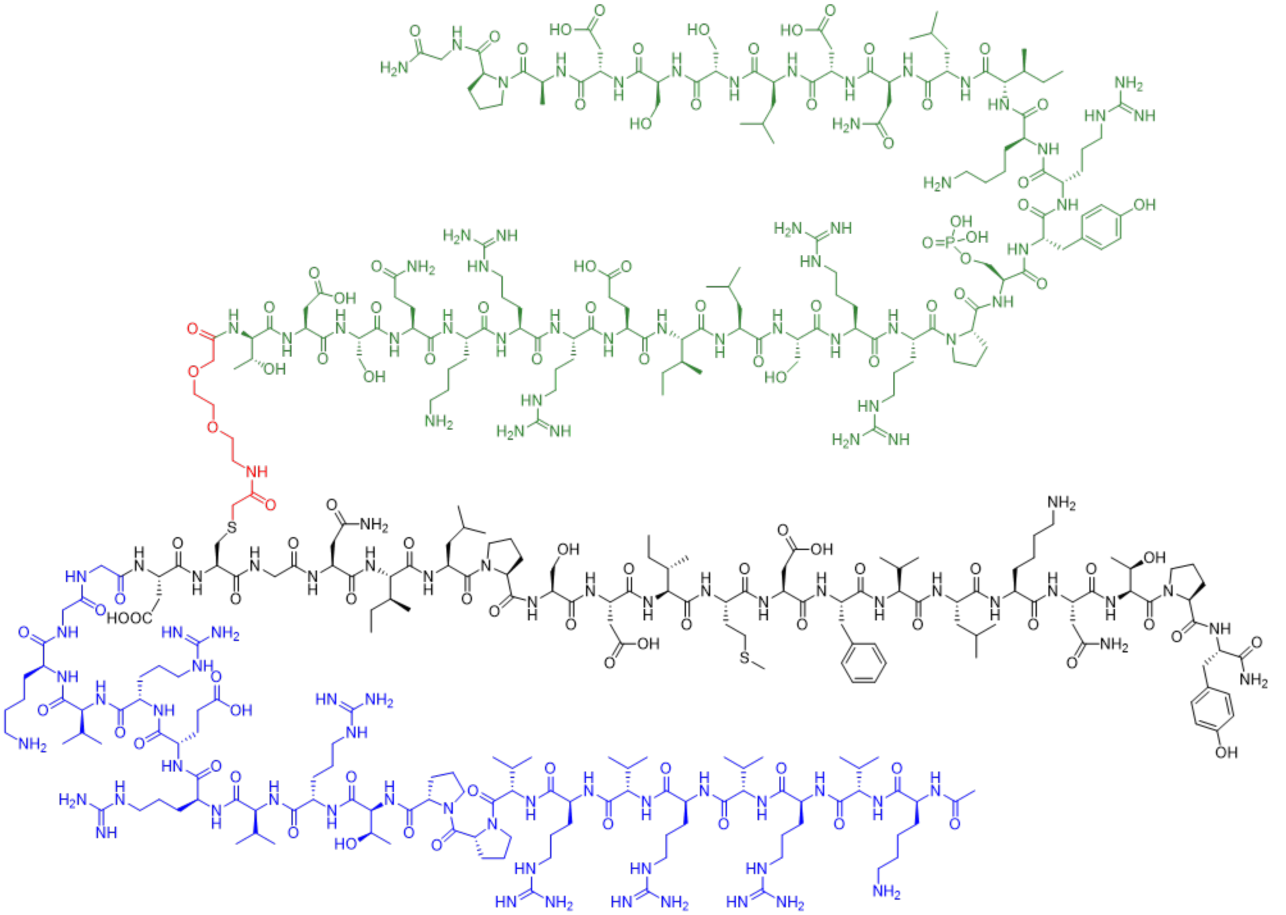

B

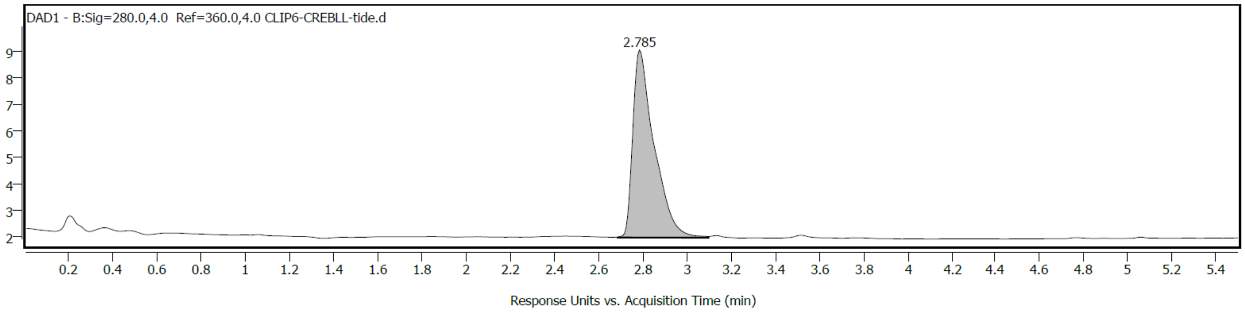
DAD1 - B:Sig=280.0,4.0 Ref=360.0,4.0 CLIP6-CREBLL-tide.d
2.785
9
8
7
6
5
4
3
2
0.2
0.4
0.6
0.8
1
1.2
1.4
1.6
1.8
2
2.2
2.4
2.6
2.8
3
3.2
3.4
3.6
3.8
4
4.2
4.4
4.6
4.8
5
5.2
5.4
Response Units vs. Acquisition Time (min)

C

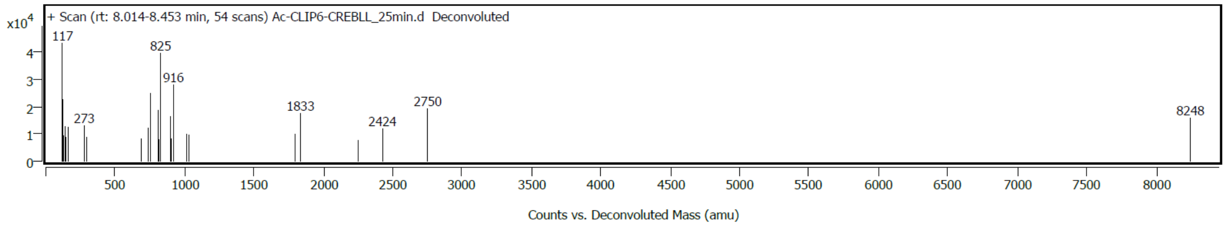


**Figure S19.** Characterization of CLIP6-CREBLL-tide. (A) Chemical structure of the peptide. (B) Analytical trace of CLIP6-CREBLL-tide monitored at 280 nm wavelength. (C) Deconvolution spectra of CLIP6-CREBLL-tide using LC-MS qTOF Mass Spectrometry.

A

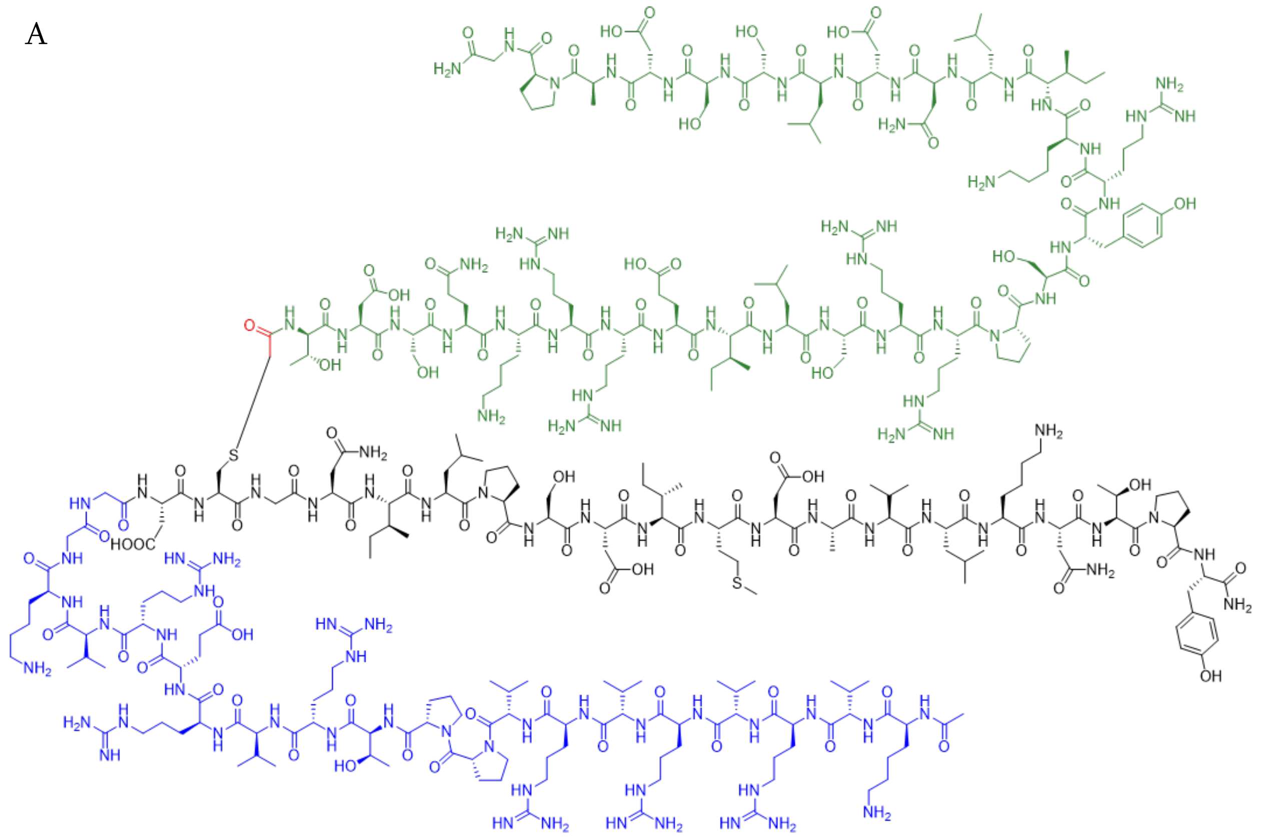

B

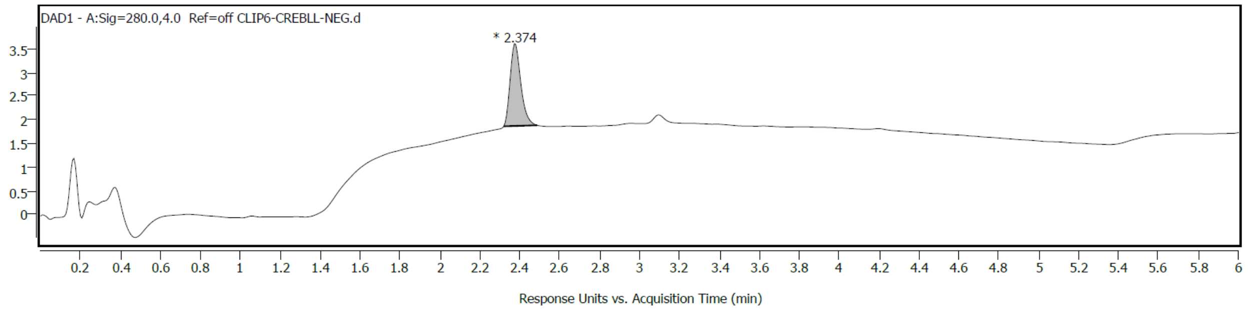
DAD1 - A:Sig=280.0,4.0 Ref=off CLIP6-CREBLL-NEG.d
* 2.374
Response Units vs. Acquisition Time (min)

A

C

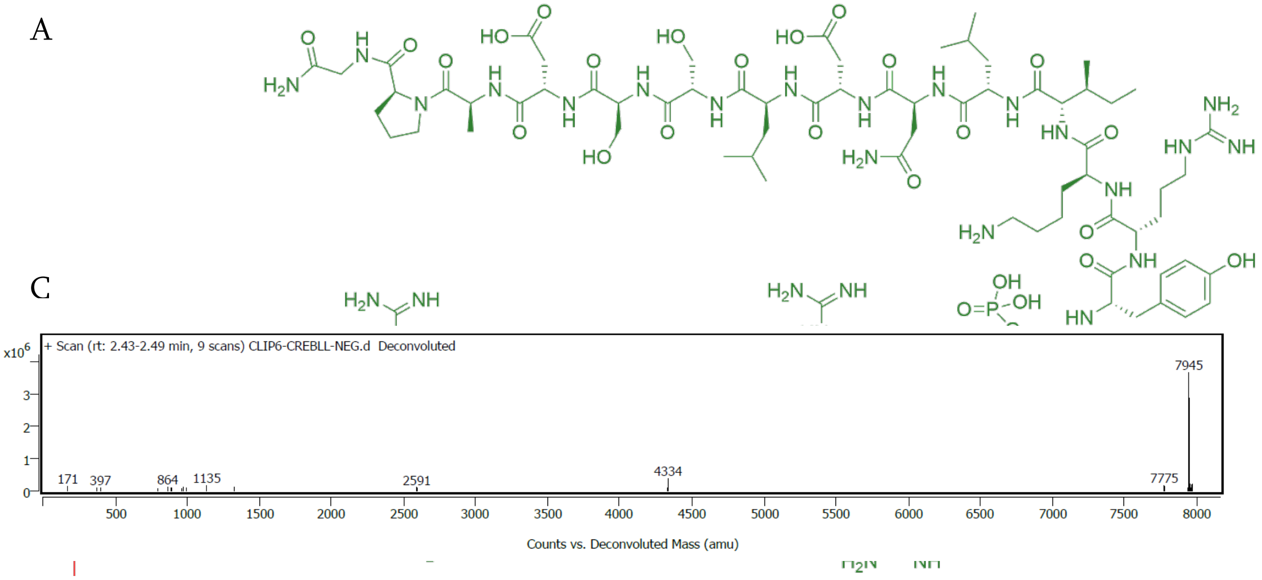


**Figure S20.** Characterization of CLIP6-CREBLL-NEG. (A) Chemical structure of the peptide. (B) Analytical trace of CLIP6-CREBLL-NEG monitored at 280 nm wavelength. (C) Deconvolution spectra of CLIP6-CREBLL-NEG using LC-MS qTOF Mass Spectrometry.

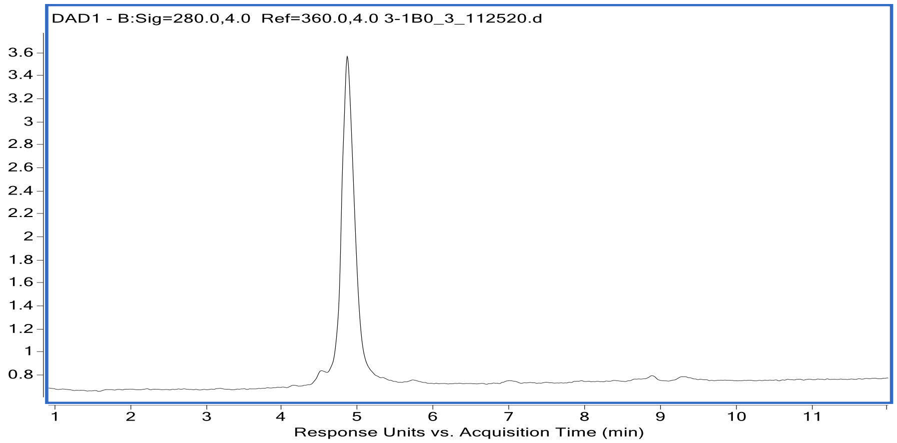


C

A

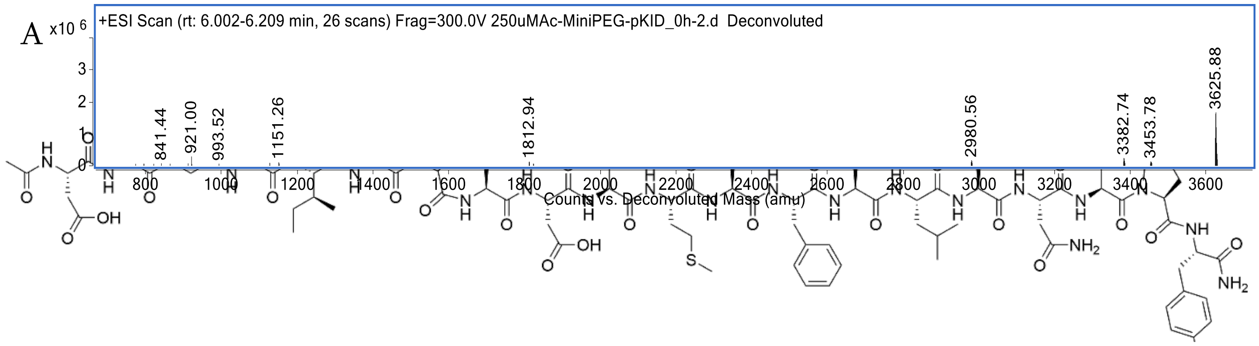


**Figure S21.** Characterization of AEEAc-pKID. (A) Chemical structure of the peptide. (B) Analytical trace of AEEAc-pKID monitored at 280 nm wavelength. (C) Deconvolution spectra of AEEAc-pKID using LC-MS qTOF Mass Spectrometry.

**Figure S22**. Characterization of MLL2840-2858. (A) Chemical structure of the peptide. (B) Analytical trace of MLL2840-2858 monitored at 280 nm wavelength. (C) Deconvolution spectra of MLL2840-2858 using LC-MS qTOF Mass Spectrometry

B

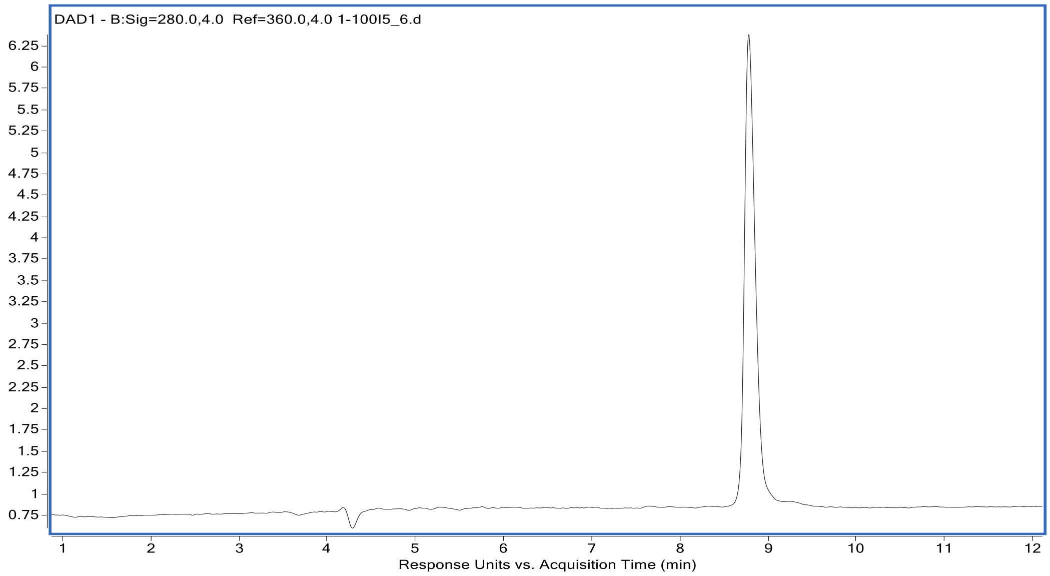


C

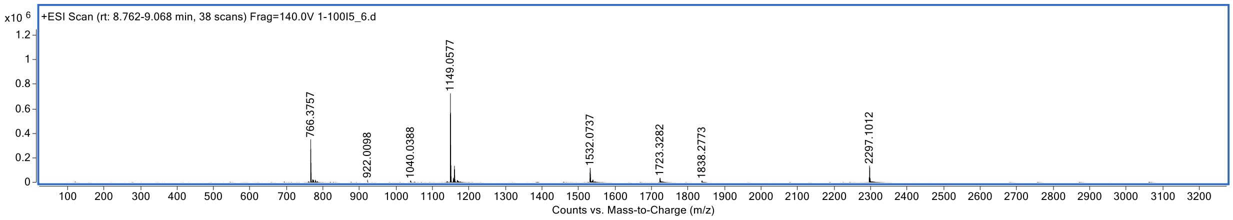

A

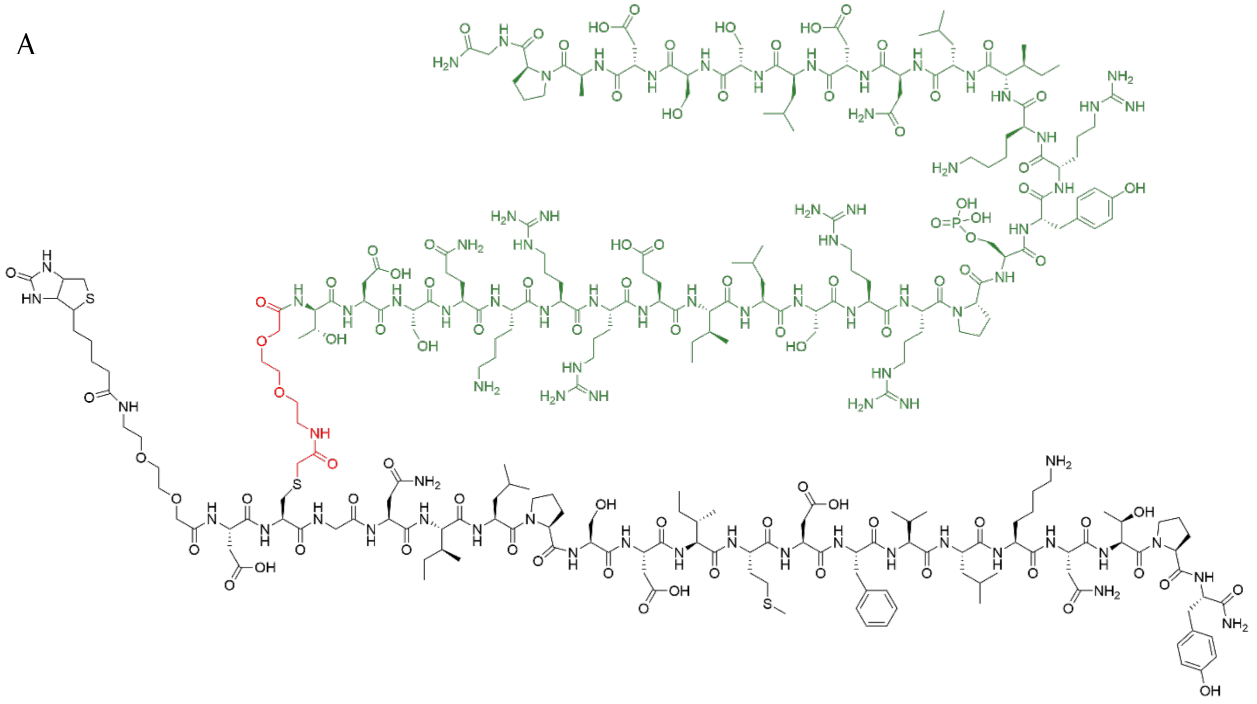

B

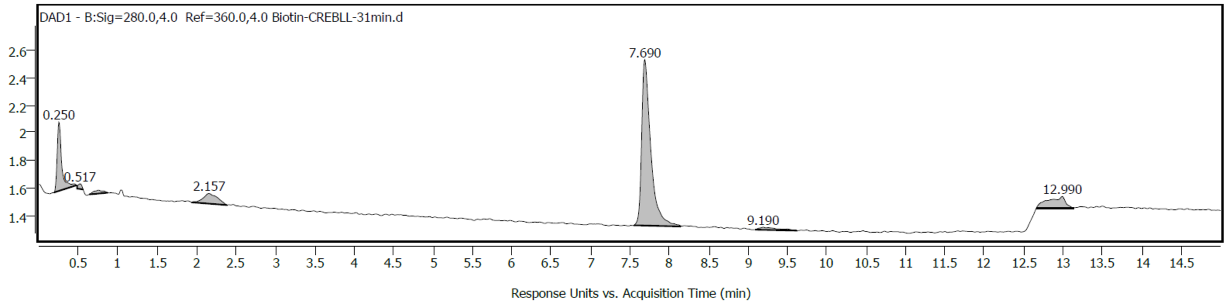


C

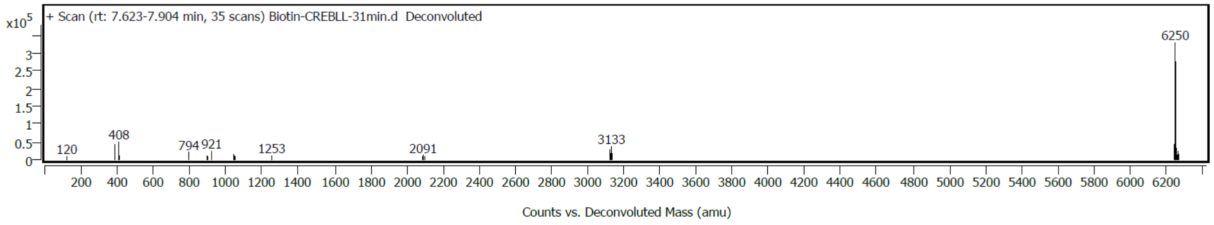


**Figure S23.** Characterization of Biotin-CREBLL-tide. (A) Chemical structure of the peptide. (B) Analytical trace of Biotin-CREBLL-tide monitored at 280 nm wavelength. (C) Deconvolution spectra of Biotin-CREBLL-tide using LC-MS qTOF Mass Spectrometry.

A

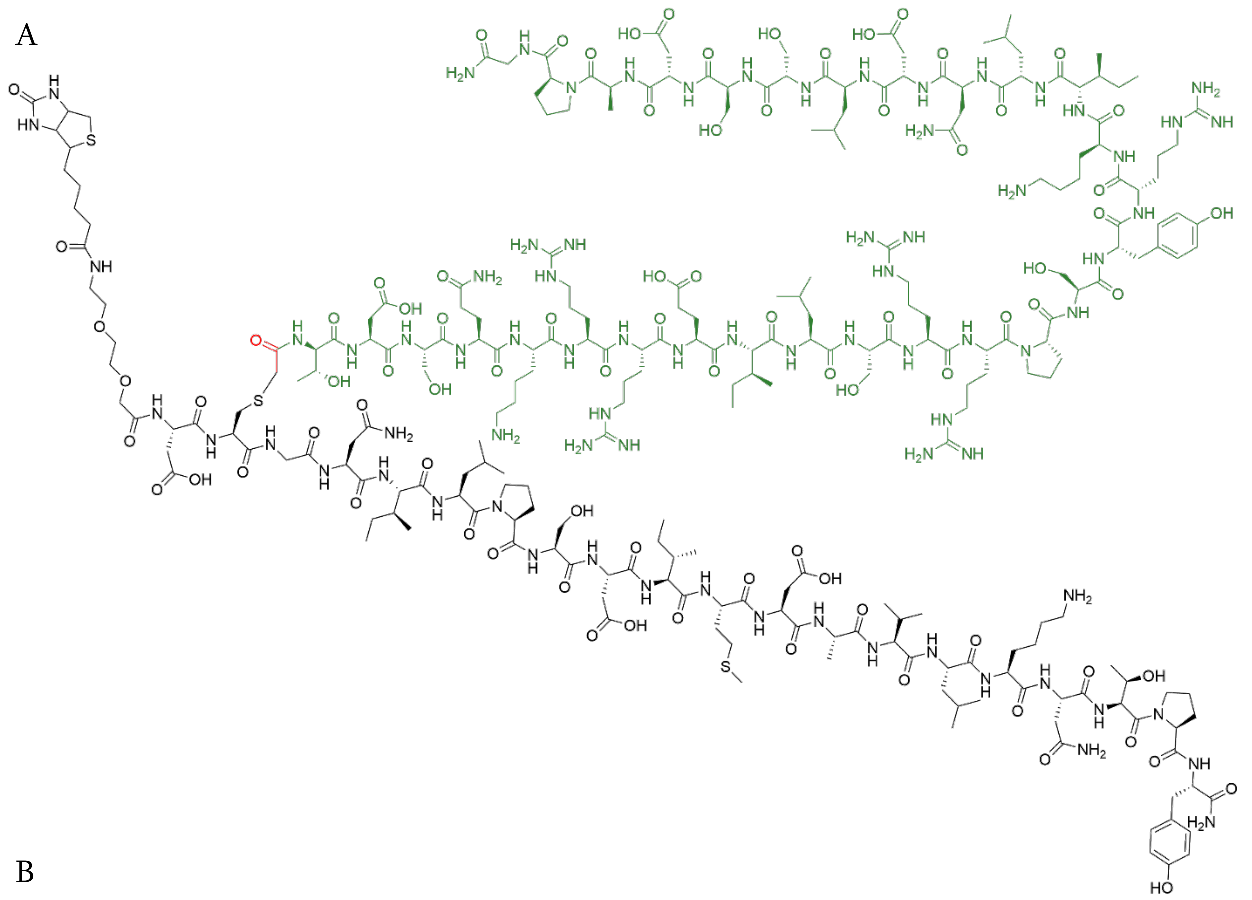

B

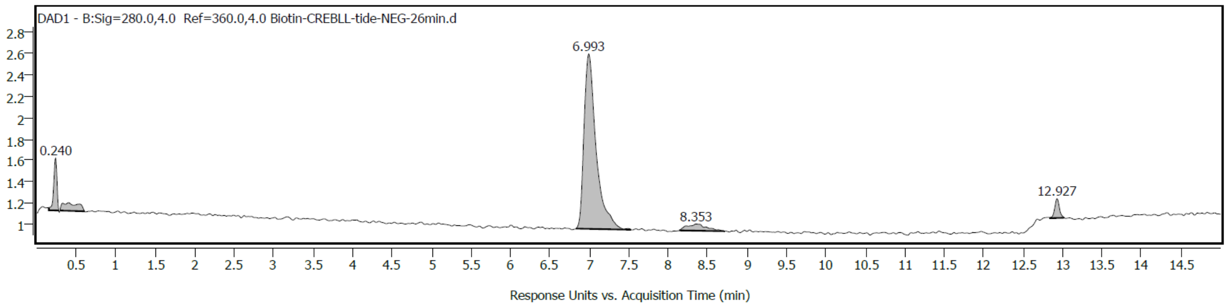
DAD1 - B:Sig=280.0,4.0 Ref=360.0,4.0 Biotin-CREBLL-tide-NEG-26min.d
0.240
6.993
8.353
12.927
Response Units vs. Acquisition Time (min)

C

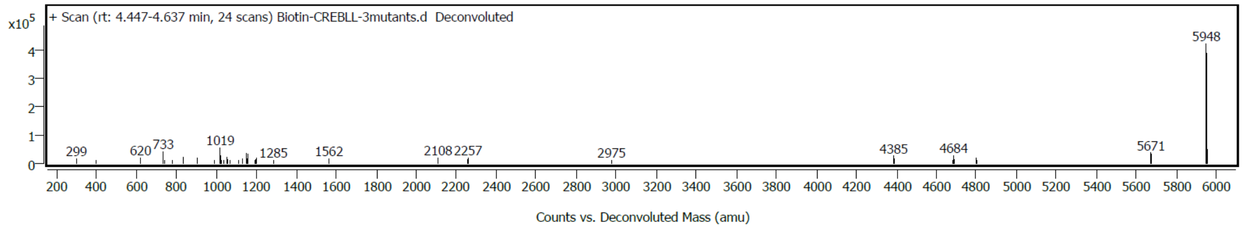


**Figure S24.** Characterization of Biotin-CREBLL-Neg. (A) Chemical structure of the peptide. (B) Analytical trace of Biotin-CREBLL-Neg monitored at 280 nm wavelength. (C) Deconvolution spectra of Biotin-CREBLL-Neg using LC-MS qTOF Mass Spectrometry.

A

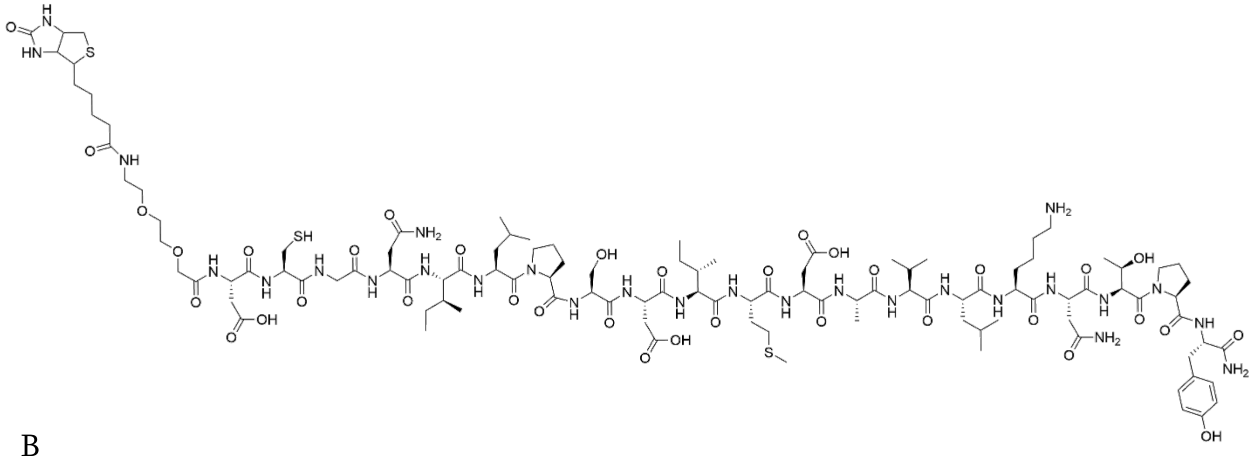

B

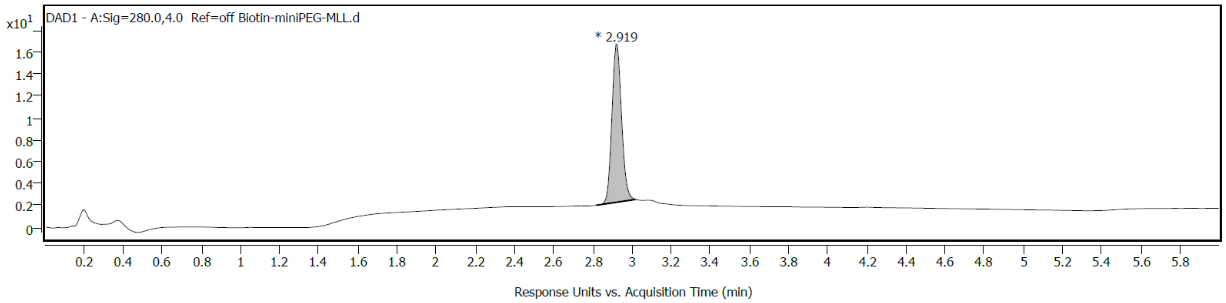


C

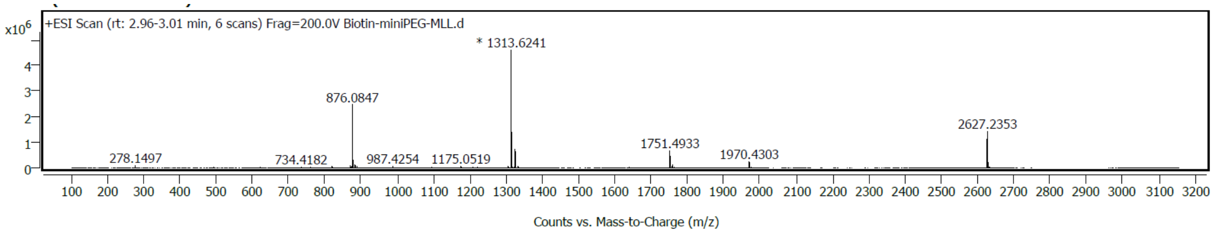


**Figure S25** Characterization of Biotin-AEEAc-MLL. (A) Chemical structure of the peptide. (B) Analytical trace of Biotin-AEEAc-MLL monitored at 280 nm wavelength. (C) Deconvolution spectra of Biotin-AEEAc-MLL using LC-MS qTOF Mass Spectrometry

A

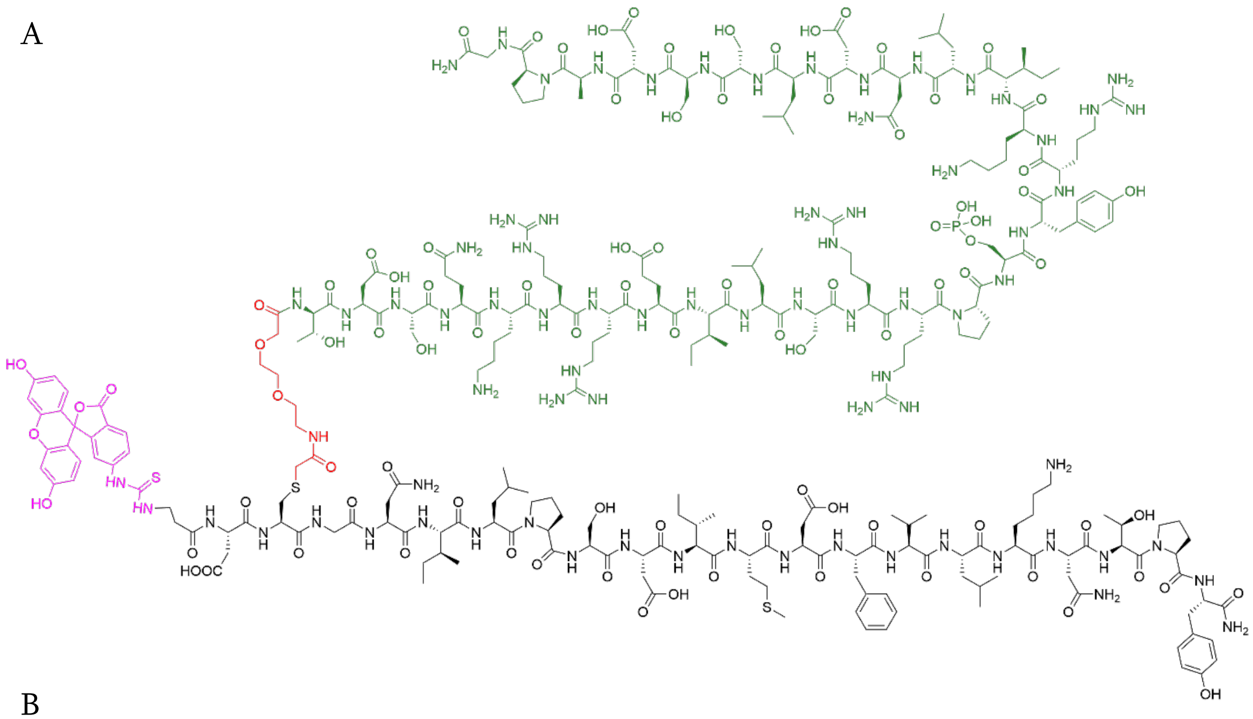

B

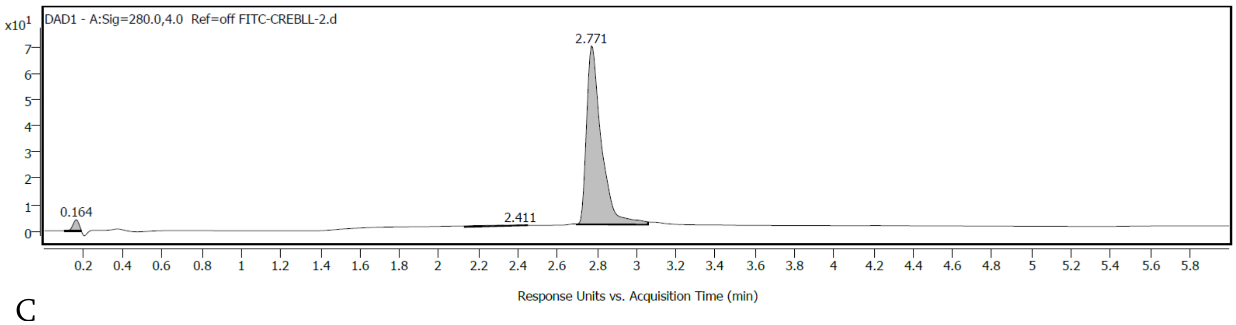


C

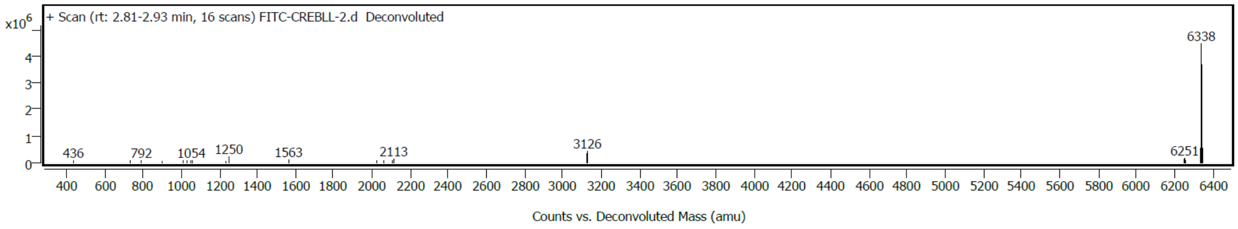


**Figure S26.** Characterization of FITC-CREBLL-tide. (A) Chemical structure of the peptide. (B) Analytical trace of FITC-CREBLL-tide monitored at 280 nm wavelength. (C) Deconvolution spectra of FITC-CREBLL-tide using LC-MS qTOF Mass Spectrometry

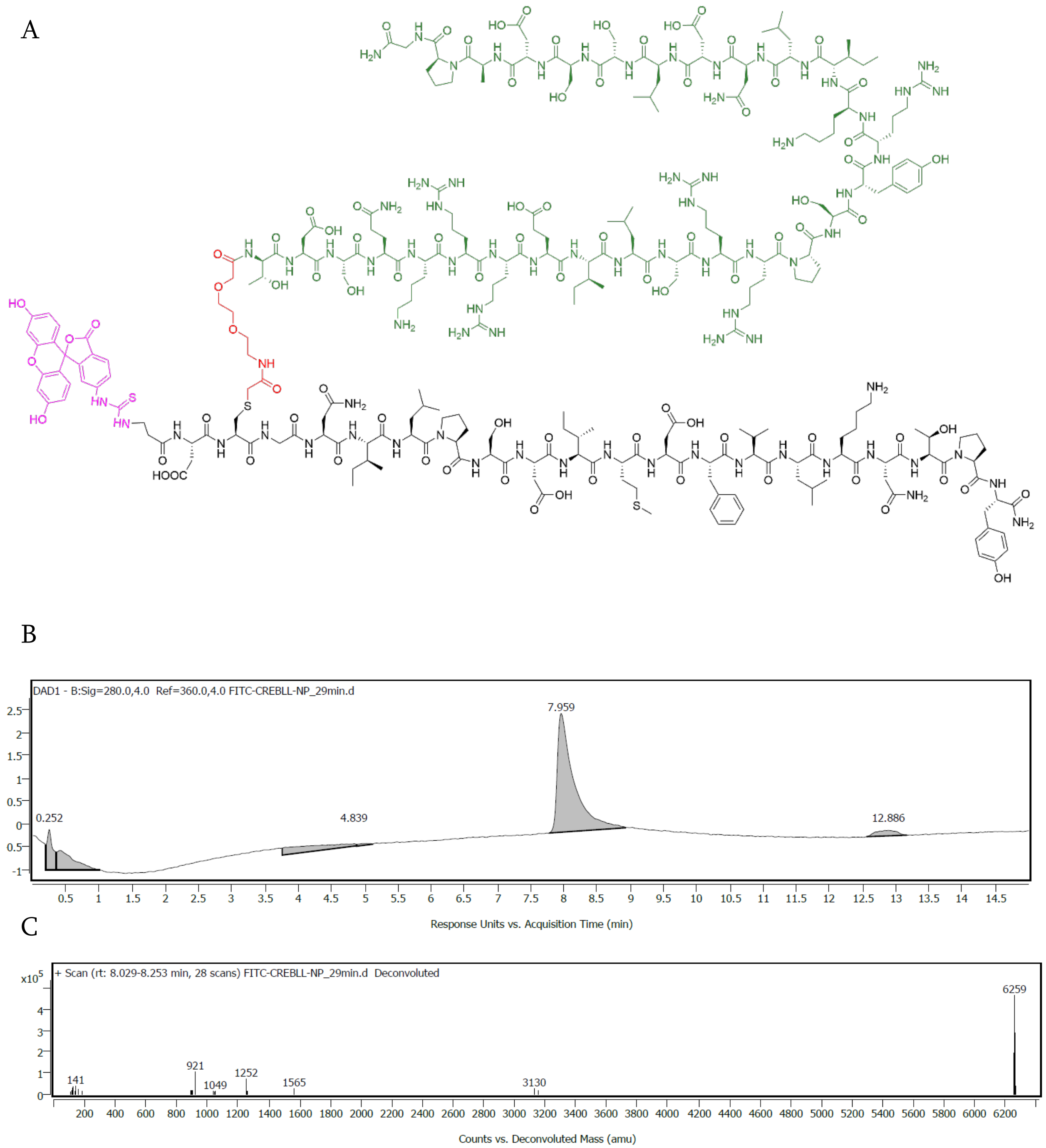


**Figure S27.** Characterization of FITC-CREBLL-tide-Nophos. (A) Chemical structure of the peptide. (B) Analytical trace of FITC-CREBLL-tide-Nophos monitored at 280 nm wavelength. (C) Deconvolution spectra of FITC-CREBLL-tide-Nophos using LC-MS qTOF Mass Spectrometry.

A

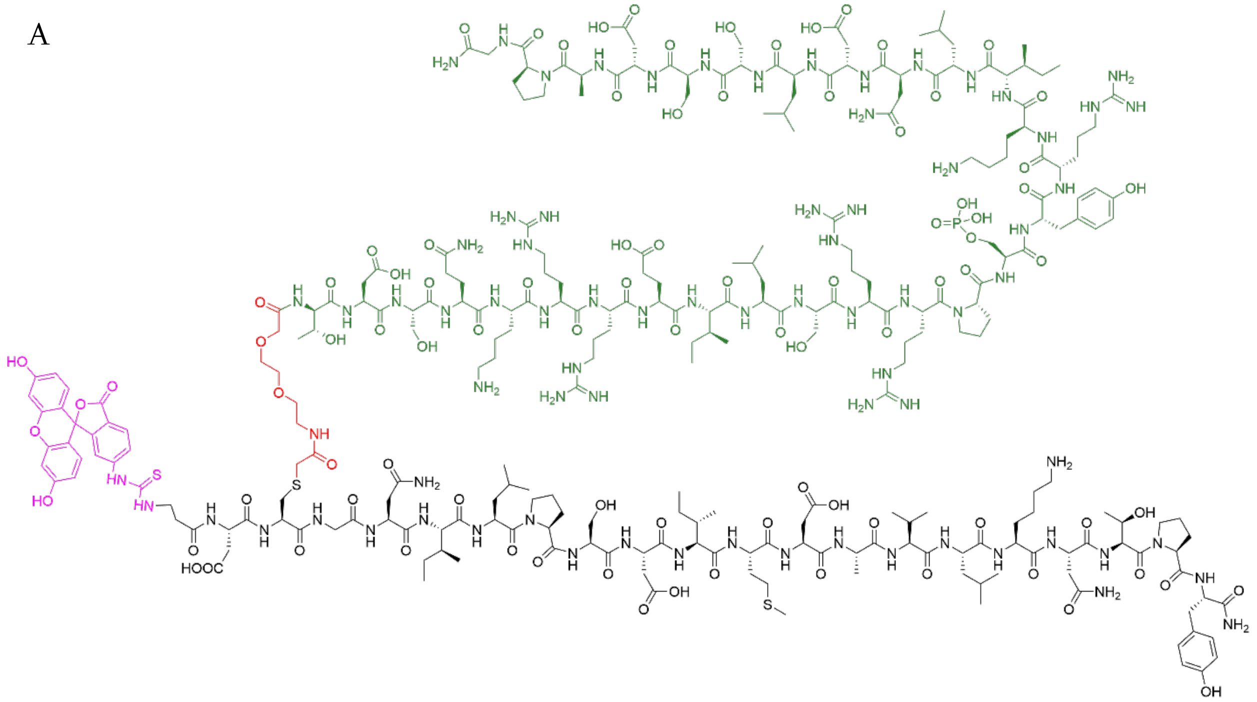

B

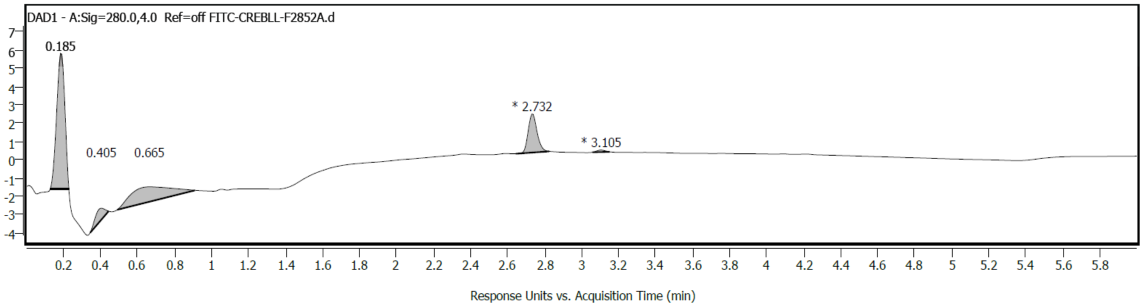


C

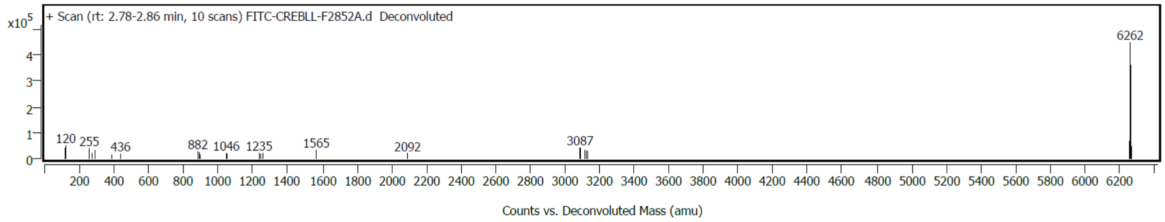


**Figure S28**. Characterization of FITC-CREBLL-tide-F2852A. (A) Chemical structure of the peptide. (B) Analytical trace of FITC-CREBLL-tide-F2852A monitored at 280 nm wavelength. (C) Deconvolution spectra of FITC-CREBLL-tide-F2852A using LC-MS qTOF Mass Spectrometry.

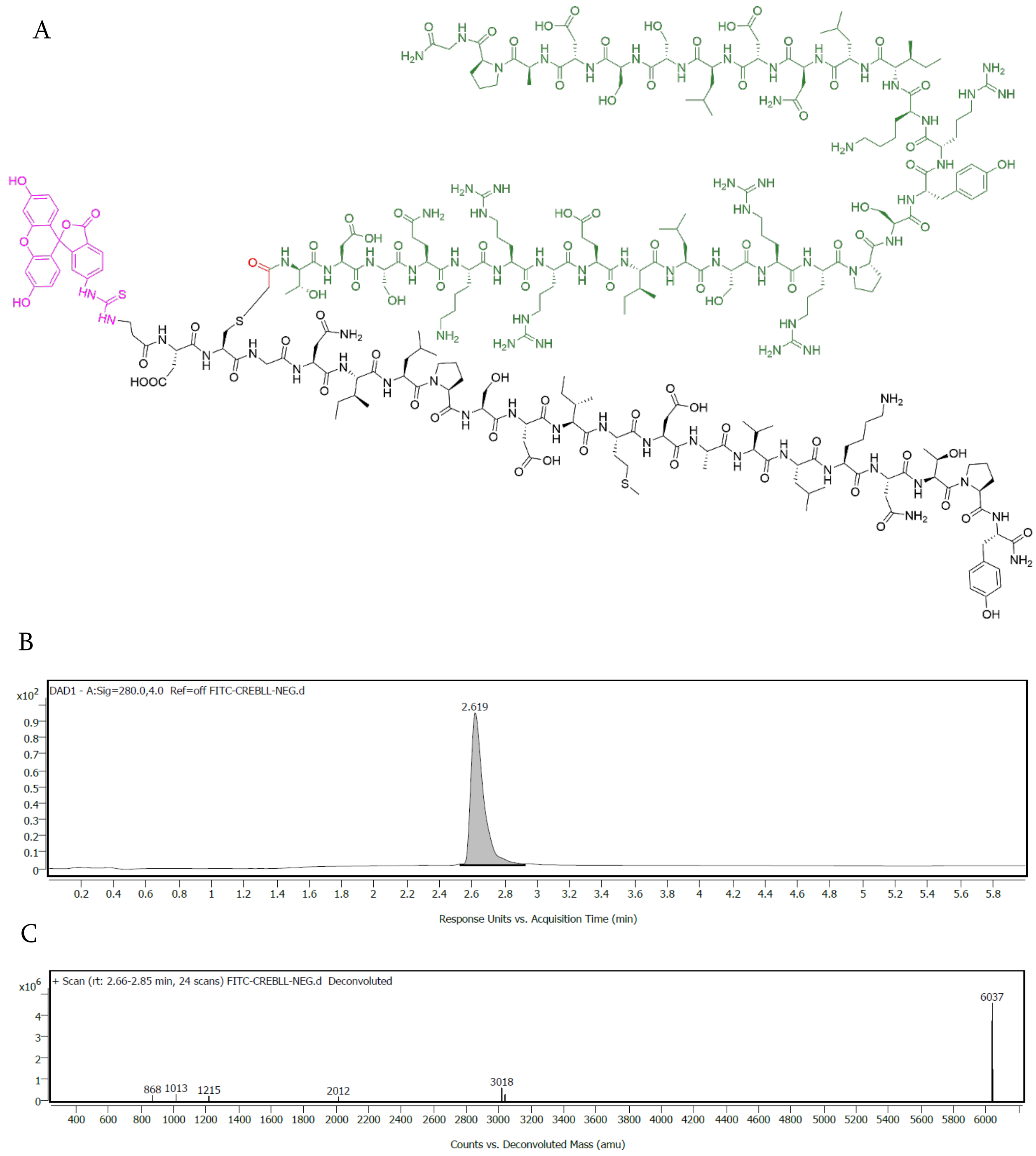


**Figure S29.** Characterization of FITC-CREBLL-tide-Neg. (A) Chemical structure of the peptide. (B) Analytical trace of FITC-CREBLL-tide-Neg monitored at 280 nm wavelength. (C) Deconvolution spectra of FITC-CREBLL-tide-Neg using LC-MS qTOF Mass Spectrometry.

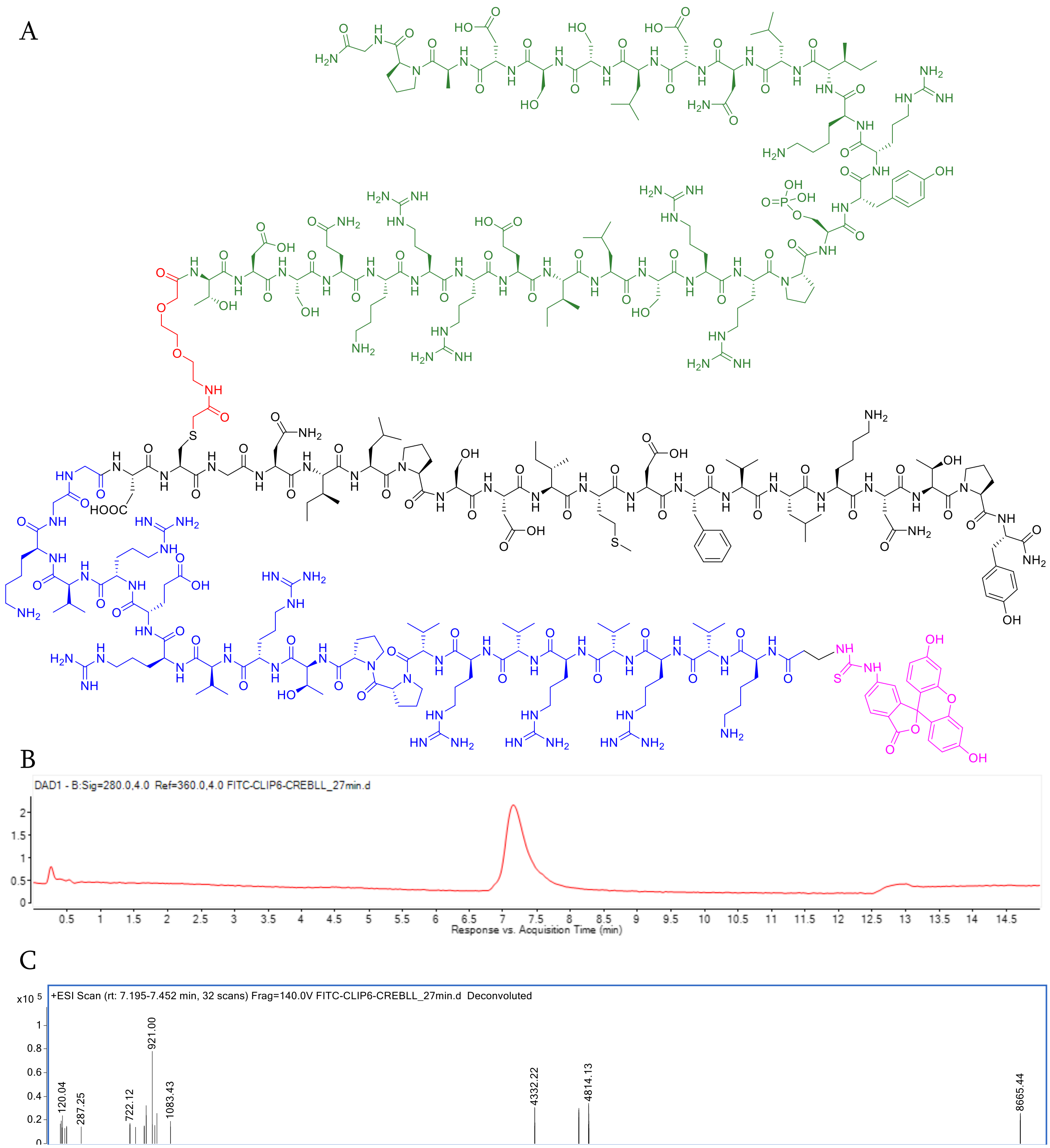


**Figure S30.** Characterization of FITC-CLIP6-CREBLL-tide. (A) Chemical structure of the peptide. (B) Analytical trace of FITC-CLIP6-CREBLL-tide Neg monitored at 280 nm wavelength. (C) Deconvolution spectra of FITC-CLIP6-CREBLL-tide using LC-MS qTOF Mass Spectrometry.

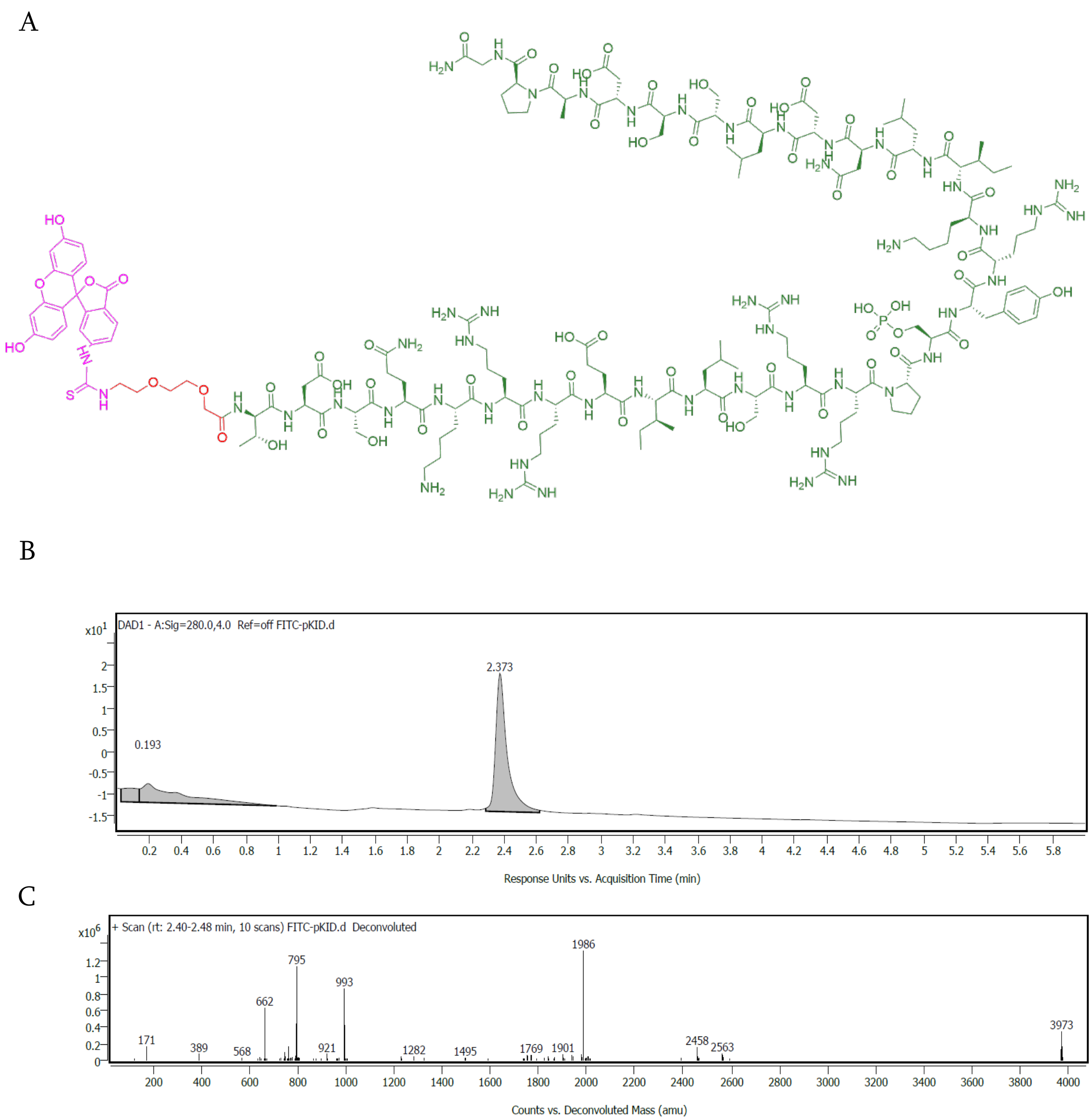


**Figure S31.** Characterization of FITC-AEEAc-pKID. (A) Chemical structure of the peptide. (B) Analytical trace of FITC-AEEAc-pKID monitored at 280 nm wavelength. (C) Deconvolution spectra of FITC-AEEAc-pKID using LC-MS qTOF Mass Spectrometry.

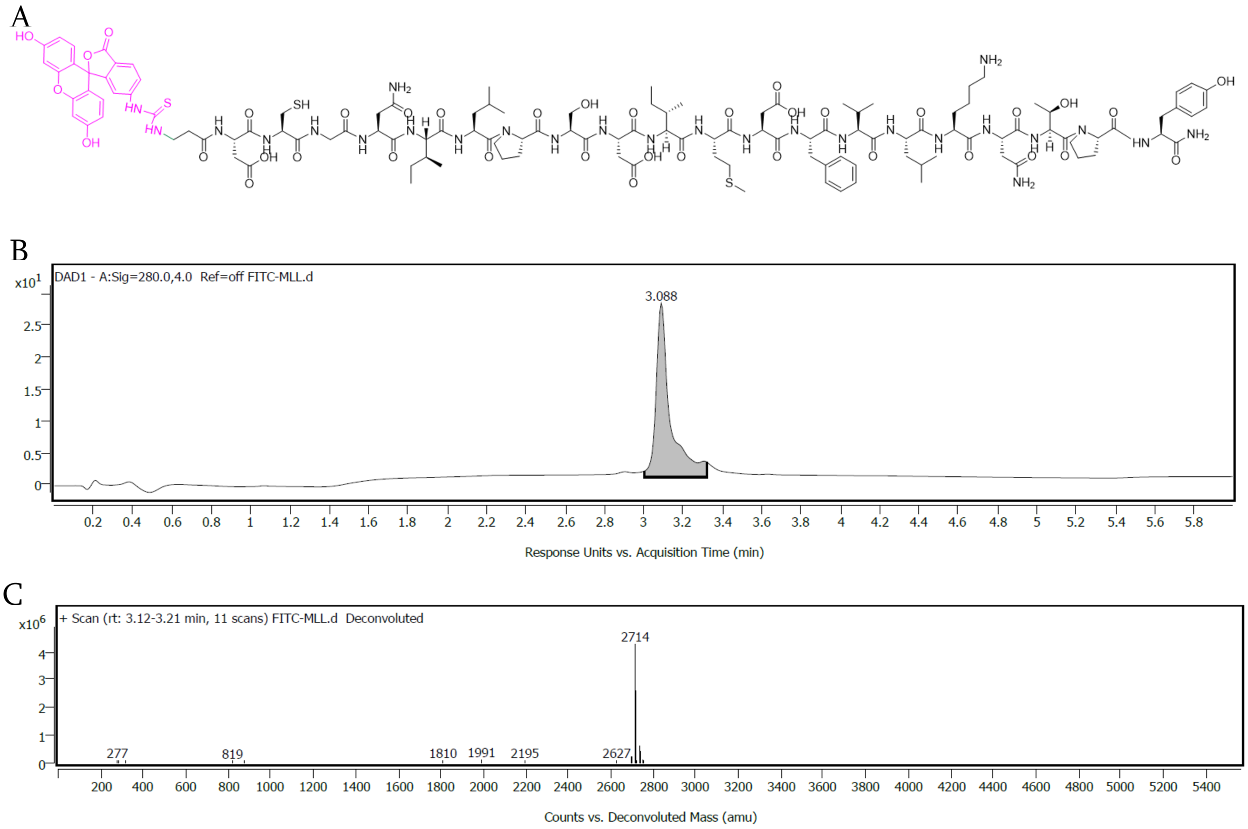


**Figure S32.** Characterization of FITC-MLL. (A) Chemical structure of the peptide. (B) Analytical trace of FITC-MLL monitored at 280 nm wavelength. (C) Deconvolution spectra of FITC-MLL using LC-MS qTOF Mass Spectrometry.

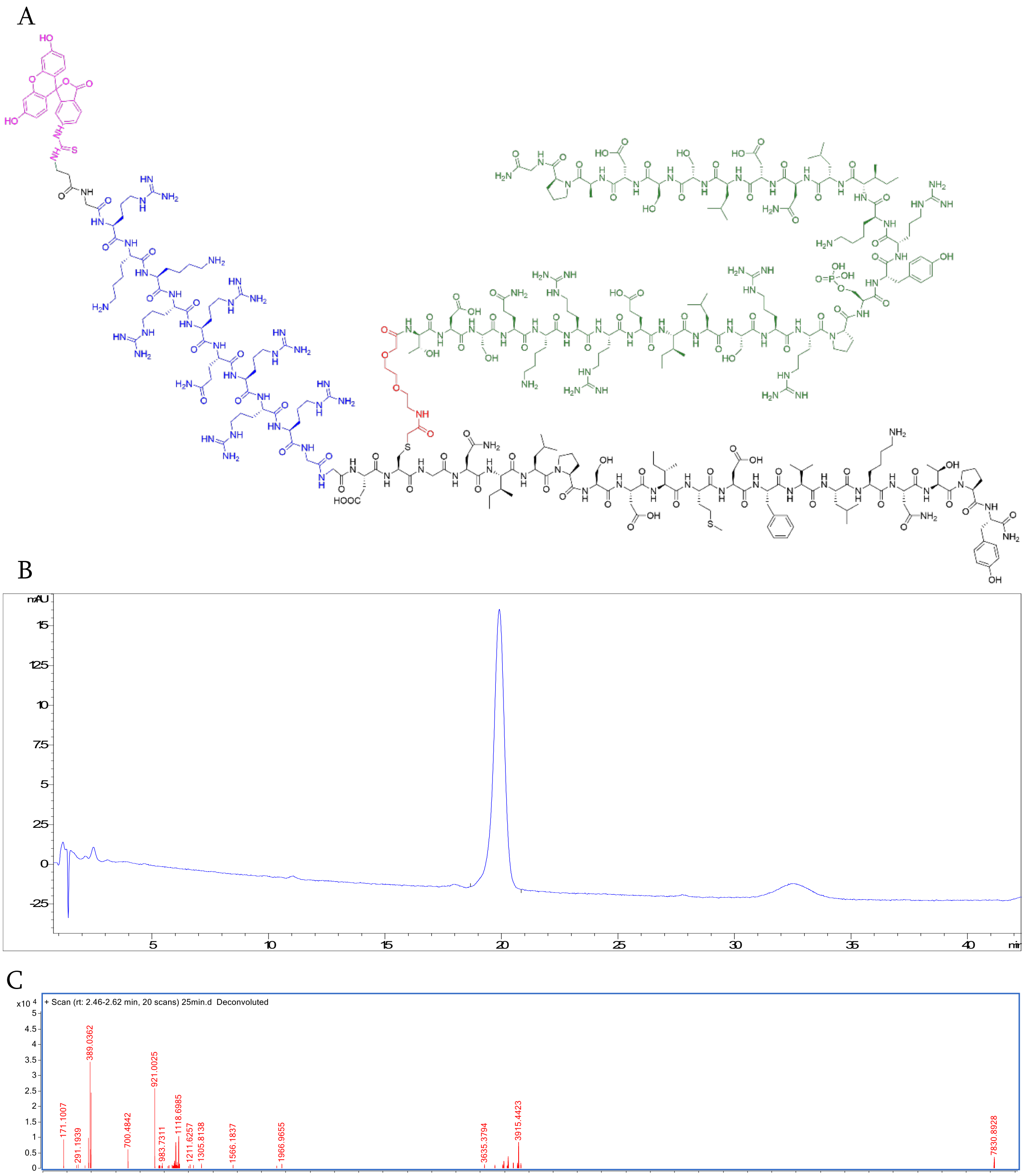


**Figure S33.** Characterization of FITC-TAT-CREBLL-tide. (A) Chemical structure of the peptide. (B) Analytical trace of FITC-TAT-CREBLL-tide monitored at 280 nm wavelength. (C) Deconvolution spectra of FITC-TAT-CREBLL-tide using LC-MS qTOF Mass Spectrometry.

14. Depmap. *https://depmap.org/portal.*
